# Structure-dependent spontaneous calcium dynamics in cultured neuronal networks on microcontact-printed substrates

**DOI:** 10.1101/2023.02.08.527409

**Authors:** Zhe Chen, Tao Sun, Zihou Wei, Xie Chen, Huaping Wang, Qiang Huang, Shingo Shimoda, Toshio Fukuda, Qing Shi

**Affiliations:** Beijing Advanced Innovation Center for Intelligent Robots and Systems, and Key Laboratory of Biomimetic Robots and Systems, Ministry of Education, Beijing Institute of Technology, Beijing 100081, China; School of Medical Technology, Beijing Institute of Technology, Beijing 100081, China; Key Laboratory of Biomimetic Robots and Systems, Ministry of Education, Beijing Institute of Technology, Beijing 100081, China; RIKEN Center for Brain Science, TOYOTA Collaboration Center, RIKEN, Nagoya, Aichi 4630003, Japan

## Abstract

Microcontact printing (*μ*CP) is widely used in neuroscience research. However, *μ*CP yields reduced cell-substrate adhesion compared with directly coating cell adhesion molecules. Here, we demonstrate that the reduced cell-substrate adhesion caused by *μ*CP, high seeding density, and the local restriction would separately contribute to more aggregated (neurons closer to each other in separate clusters) neuronal networks. Calcium recordings revealed that more aggregated networks presented fewer spontaneous calcium activity patterns, and were more likely dominated by synchronized network-wide calcium oscillation (network bursts). First, on a uniform microcontact-printed substrate, densely seeded neurons were reaggregated into a Petri dish-wide network consisting of small clusters, of which the calcium dynamics were dominated by network bursts. Next, further analysis revealed this dominance was maintained since its appearance, and the initiation and propagation of bursts in the small-cluster network shared a similar mechanism with that of homogeneous networks. Then, sparsely seeded neurons formed several networks with different aggregation degrees, in which the less clustered ones presented abundant time-varying subnetwork burst patterns. Finally, by printing locally restricted patterns, highly clustered networks formed, where dominant network bursts reappeared. These findings demonstrate the existence of structure-dependent spontaneous calcium dynamics in cultured networks on microcontact-printed substrates, which provide important insights into designing cultured networks by using *μ*CP, and into deciphering the onset and evolution of network bursts in developmental nerve systems.

## I. INTRODUCTION

With the advancement of biofabrication technologies, engineering neuronal networks *in vitro* to recapitulate the organization and functionality of neural circuits *in vivo* has attracted increasing attention in the last two decades [1–6]. It complements computational modeling and holds the potential to elucidate the functional mechanism of neural systems at both network and cellular levels, since it allows neuroscientists to custom design neural circuits biologically [7–11]. Neuronal networks mimicking brain circuits could be built because dissociated primary neurons could form functional synapses even on non-physiological surfaces [12–18].

Among the bio-interface technologies, microcontact printing (*μ*CP) has gained popularity in reconstructing neuronal networks *in vitro*, owing to its ease of use, high throughput, versatility and reusability [19–23]. However, *μ*CP produces a substantially thinner cell adhesion molecules (CAM) layer because of the limited adsorption time and the protein transfer process, compared with directly coating cell adhesion molecules [24]. Such a thinner CAM layer reduces cell-substrate adhesion compared with CAM-coated substrates. Neurons formed a homogeneous network on substrates with high cell-substrate adhesion, e.g., a carpet of glia [25] or a thick layer of coated CAM [26], and reaggregated to form a clustered network on substrates with low cell-substrate adhesion, e.g., untreated glass substrates [27] or ploy-D-lysine precoated polystyrene (PS) surface [28]. Research have shown that, on CAM-coated substrates, the neuron-substrate adhesion [27, 29], seeding density [30, 31], and the local topological restriction [3, 8, 32] will impact not only the morphology and structure but also the spontaneous calcium dynamics of cultured neuronal networks, which reflects the intrinsic functional connectivity within certain networks [33]. However, on microcontact-printed substrates, the combined influence of these factors on network structure and calcium dynamics remains to be investigated.

In this work, we combined *μ*CP with calcium fluorescence imaging to explore the impact of reduced cell-substrate adhesion caused by *μ*CP, seeding density and local restriction on the network structure and spontaneous calcium activity of cultured neuronal networks, as shown in Fig.1. First, on a uniform microcontact-printed substrate, a Petri dish-wide neuronal network consisting of interconnected small clusters was obtained. In this small-cluster network, network-wide bursts dominated the calcium dynamics. Then, we reduced the cell seeding density, and several networks with different clustering degrees formed. In the less clustered ones, abundant time-varying subnetwork bursts appeared, which suggested the dependence of spontaneous calcium dynamics on network structure. To examine this hypothesis, we finally leveraged patterned *μ*CP to apply local restriction, where highly clustered networks formed, of which the calcium dynamics was again dominated by network bursts.

**FIG. 1.**
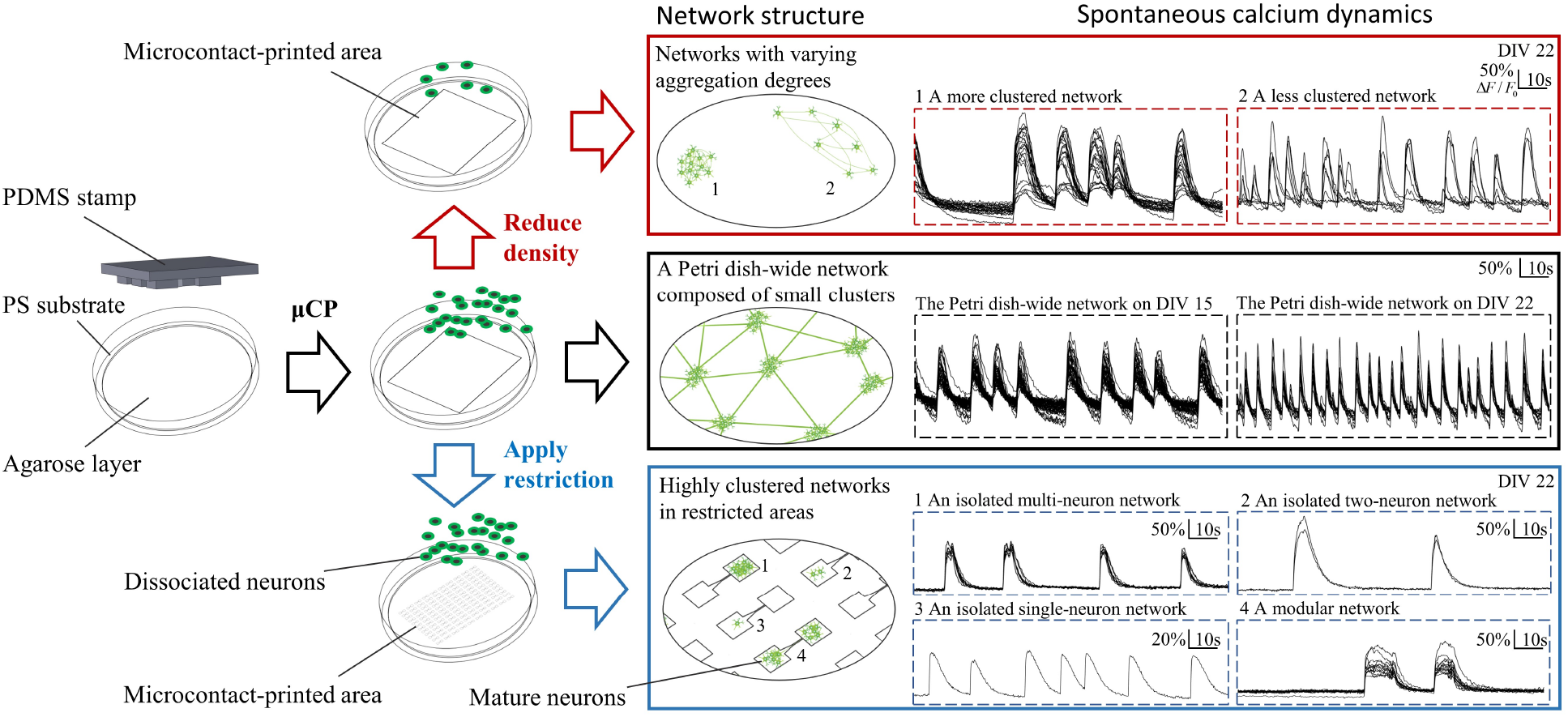
Cultured neuronal networks with different structures formed on microcontact-printed substrates by varying printed patterns and seeding density, and presented organization-related calcium activities. Middle panels: with a uniform pattern, a large-scale network composed of small clusters formed, with calcium activities characterized by network bursts. Top panels: by reducing seeding density, networks with varying clustering degrees formed, among which the less-clustered one presented time-varying subnetwork bursts. Bottom panels: by confining neurons in small squares, we obtained highly clustered networks, where dominant network bursts reappeared.

## II. RESULTS

### A. Calcium dynamics in small-cluster networks

First, we printed protein ink on an agarose-treated polystyrene (PS) substrate using an unpatterned polydimethylsiloxane (PDMS) stamp to explore how μCP-reduced cell-substrate adhesion will influence neuronal network structure and function (see supplementary material for details) [Figs. S1 and S2]. We found that, on such a substrate, isolated neurons with high seeding density tended to reaggregate into small clusters [Figs. 2(a),(b)]. Further inspection revealed that the small clusters connected with neighboring ones, forming a single small-cluster large-scale network [Fig. S4]. The maturation of such a network was also delayed, compared with homogeneous [34–36] and large-cluster[27, 29] networks, both of which demonstrated stable synchronized calcium oscillation as early as day 5 after plating. In the small-cluster network, such a calcium activity was not observed until day 15. On day 9, only sporadic cluster-level or subcluster bursts consisting of fewer than 6 neurons were observed under calcium fluorescence recording [Fig. S3]. However, on day 15, a functional network formed, which was evidenced by spontaneous network-wide synchronized calcium oscillation (or “network bursts”) [Fig. 2(c)] (see supplementary movie S1), a typical phenomenon in homogeneous cultured networks reported by other researchers [20, 37]. Very few sporadic subnetwork or neuronal bursts were observed. The correlation coefficient (CC) within the network was close to 1, which indicated the high functional connectivity within the network. Overlapped traces of different neurons also revealed the high synchronicity between different parts of the network [Fig. S5(A)]. On day 22, the synchronicity remained unchanged, with the mean CC slightly reduced from 0.90 to 0.88 [Fig. 2(d)]. The CC remained high after day 15; consistently with much research on homogeneous networks, network bursts remained dominant from their appearance [36, 38].

**FIG. 2.**
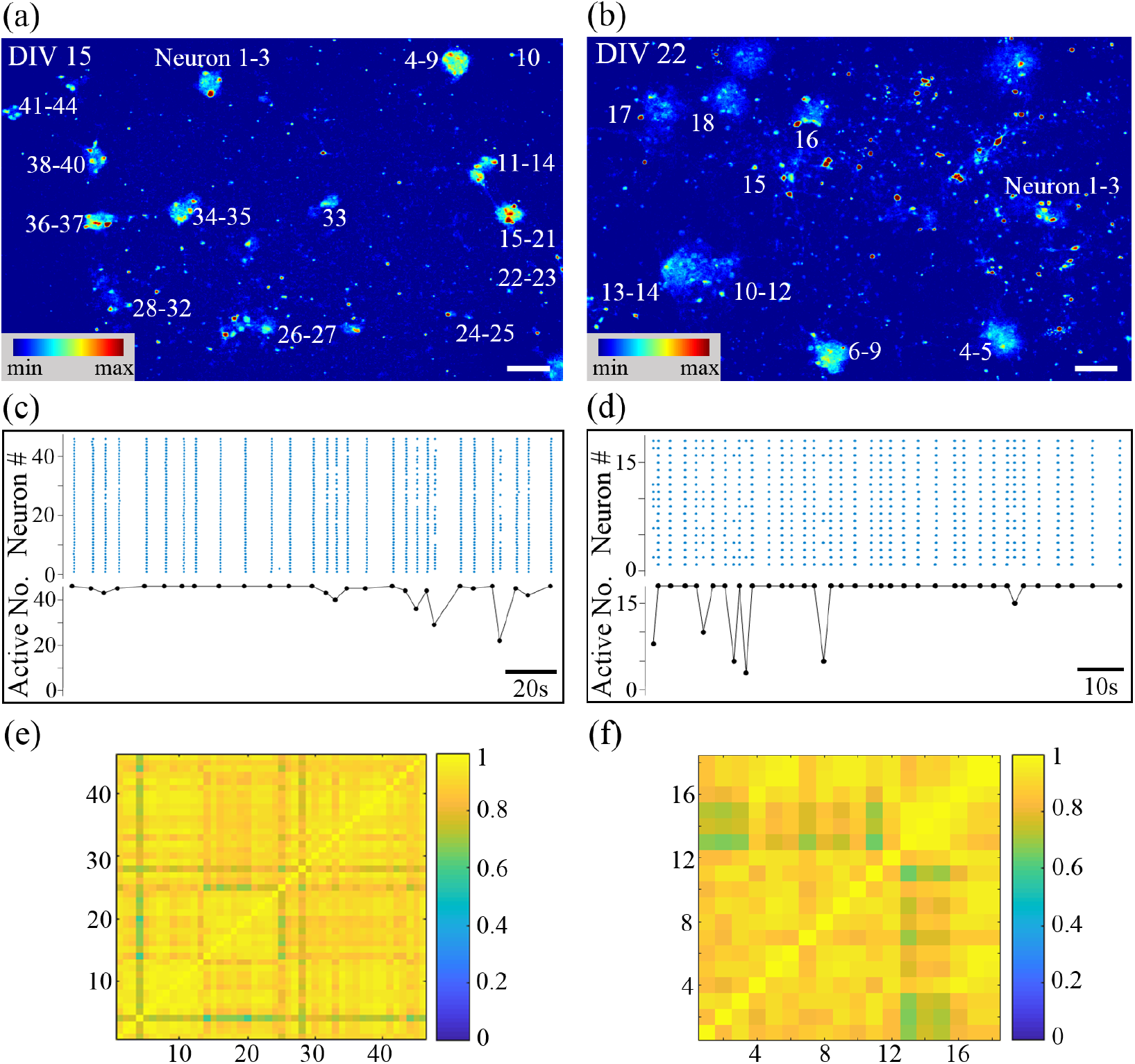
Calcium dynamics of a small-cluster network on day 15 and day 22 are both dominated by synchronized network bursts. False-color fluorescence micrographs of the network on day 15 (a) and on day 22 (b). (c) and (d) represent day 15 and day 22, respectively. Top panel: raster plot of neuronal bursts; bottom panel: number of active neurons in one subnetwork burst versus time. The matrix of CC on day 15 (e) and on day 22 (f). Scale bar: 100 *μm*.

Next, we developed an automated data processing framework to quantitatively analyze the burst waveform variation of the small-cluster network from day 15 to day 22. This framework calculated a number of important high-order characterizations by feeding the time-varying fluorescence traces of selected regions of interest (ROIs) (see Methods for details) [39]. As shown in Fig. 3, from day 15 to day 22, in the small-cluster network, the burst frequency increased from 0.15 Hz to 0.32 Hz, the burst duration reduced from 0.73 s to 0.38 s, the IBI was downregulated from 6.68 s to 3.07 s, and the calcium decay time constant was also diminished from 2.01 s to 0.69 s, consistently with the intuitive observation in Fig. S5. With network maturation, more synapses formed, the internal network information flow was reinforced, and a full-blown burst would be reached more quickly with the stronger intra-network communication, i.e., less spikes induced in each neuronal burst, explaining the shorter burst duration [35, 36, 38, 40]. Consequently, the inner peak calcium concentration reduced and the quiescent period (IBI) shortened. Therefore, the calcium ejection constant reduced and the burst frequency increased. Moreover, detailed analysis of the distribution of IBI in such networks on the two distinct time points revealed they could both be fitted well by Gaussian distributions, with *R*^2^ > 0.93 each time. It is highly possible that the network bursts happened rhythmically, added by a Poisson process [Fig. S8]. Further analysis of the distribution of IBICV confirmed such a hypothesis [Fig. S9], consistently with other homogeneous networks reported [16, 35, 36, 41]. This suggests network bursts in homogeneous and small-cluster networks could share a similar initiation and propagation mechanism, in spite of their distinct network organization. The distribution of the burst duration in these two networks could also be fitted well by Gaussian distributions, with *R*^2^ both bigger than 0.98, revealing that the fluctuation of burst duration from its mean value was probably caused by random noise.

**FIG. 3.**
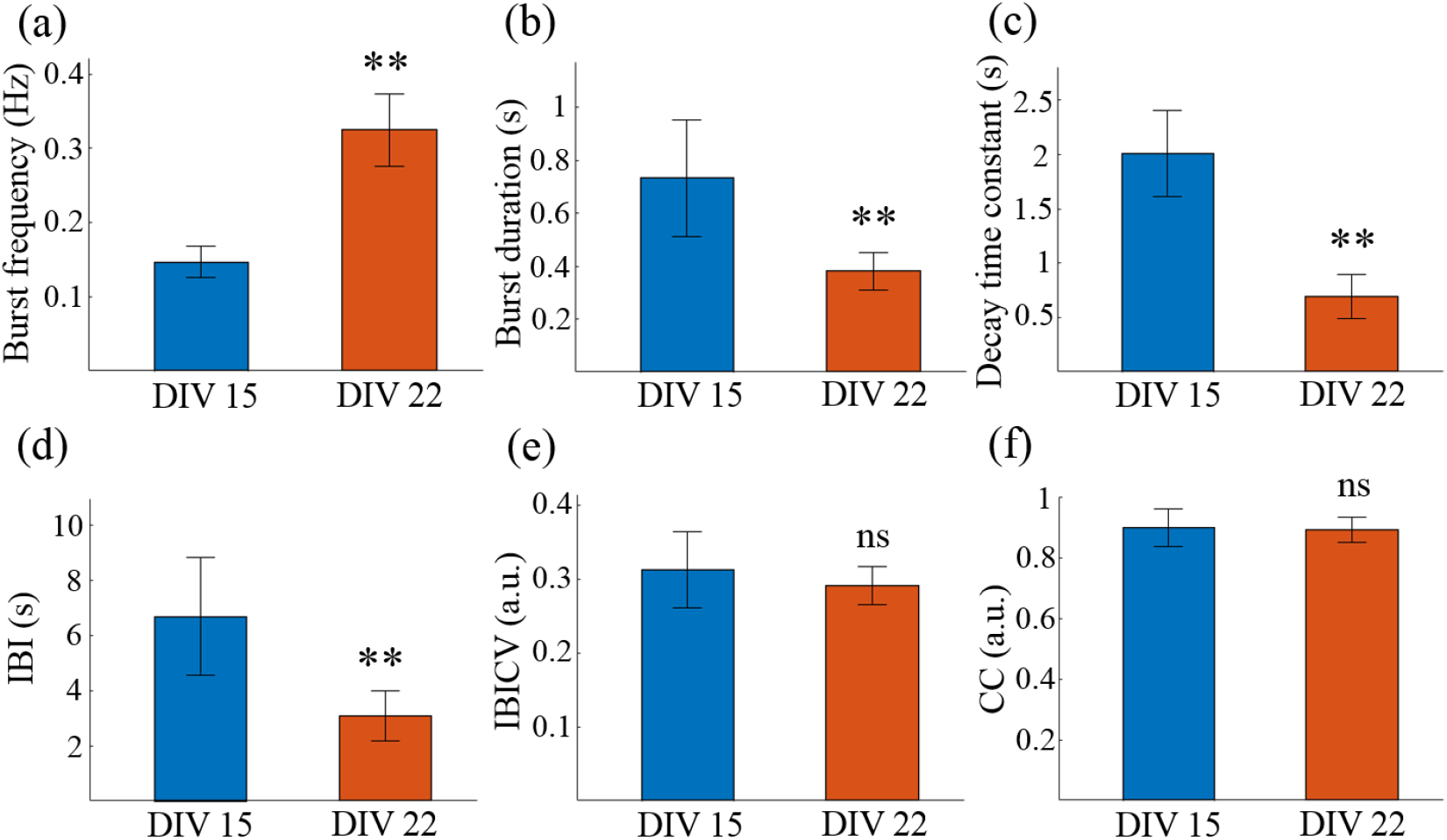
Characterization and comparison of the calcium dynamics of the small-cluster network on day 15 and on day 22. (a) Neuronal burst frequency, the average number of neuronal bursts per neuron per second. (b) Neuronal burst duration, the difference value between the onset and the end time of a neuronal burst. (c) The fitted decay time constant of the calcium attenuation phase. (d) Inter-burst interval (IBI), the interval between two consecutive neuronal bursts. (e) Coefficient of variance of IBI (IBICV), the standard deviation of the IBI distribution across neurons divided by its mean value. (f) Correlation coefficient (CC). **p < 0.01. Data are presented with mean *±* SEM.

### B. Calcium dynamics in networks of different aggregation degree

Then, to investigate the impact of diminishing interneuronal attraction on network organization, we reduced the seeding density of primary cortical neurons from 600 *mm*^−2^ to 150 *mm*^−2^, with other variables remaining unchanged. In this sparsely seeded unpatterned group, owing to the relatively long distance between neurons, neurons didn’t form clusters in the first few days after seeding. With the proliferation and spread of neuronal supporting cells, neurons sprouted neurites and built synaptic connections in the next days. Gradually, many weakly separated networks with different clustering degrees formed. They were only weakly separated because the gap between different networks could potentially be bridged with the further maturation of the culture. Such networks contained the number of neurons ranged from several to hundreds and typically colonized an area smaller than 1 *mm*^2^. In Fig. 4(a), the two highly clustered networks were so close to each other that they could not be morphologically grouped as two independent networks. However, calcium dynamics revealed that the two networks burst non-synchronously. In Fig. 4(a3), two burst patterns appear and they switch between each other randomly, indicating that the neural signal is not relayed between the two networks. Further analysis reinforced this hypothesis by showing significant distinction in CC between the two networks [Figs. 4(a2),(a4)], and higher *R*^2^ in separate networks compared to the merged one [Fig. S10]. Such a phenomenon of two neighboring networks bursting asynchronously was only reported previously for cortical networks on geographically constrained substrates [20, 32, 42–44].

**FIG. 4.**
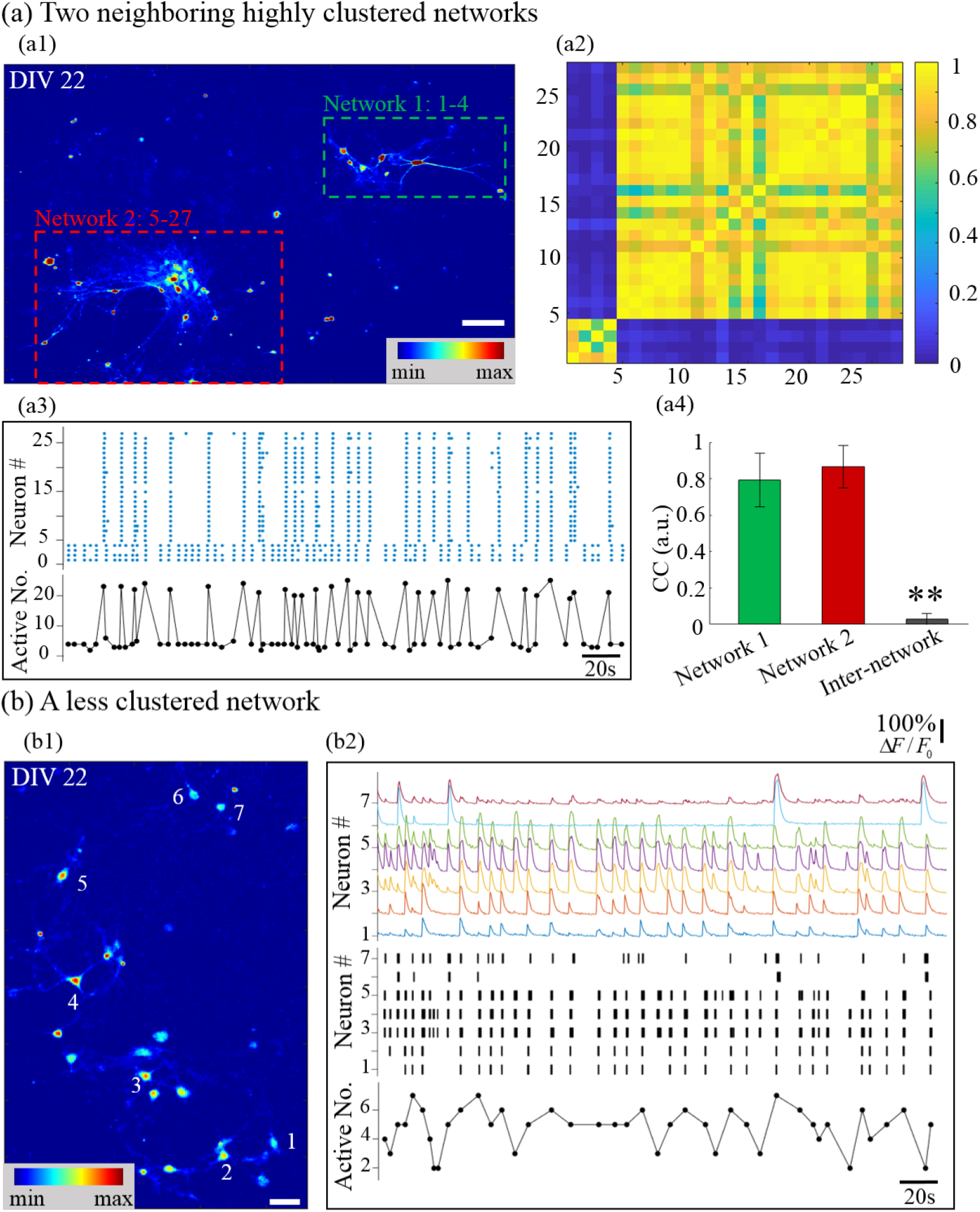
Networks with different clustering degrees present different calcium dynamics, ranging from dominant network bursts to time-varying subnetwork bursts. (a1) False-color fluorescence micrograph of two neighboring highly-clustered networks. (a2) Matrix of CC. (a3) Top panel: raster plot of neuronal bursts. (a4) CC of intra- and inter- network. (b1) False-color fluorescence micrograph of a less-clustered network consisting of seven neurons, with their locations indicated. (b2) Top panel: relative fluorescence traces of the seven neurons in (b1); middle panel: the black lines represent location of neuronal burst, with the line width coded by burst duration. Bottom panels in (a3) and (b2): active neuron number in a subnetwork burst versus time. Data are presented with mean *±* SD. **p < 0.01 against other two networks. Scale bar: (a1) 100 *μm*, (b1) 40 *μm*.

More interestingly, in other less clustered networks, time-varying subnetwork bursts appeared [Fig. 4(b)]. The seven-neuron subnetwork in Fig. 4(b) was abstracted from a weakly separated network and treated as an independent network, in which abundant time-varying subnetwork burst patterns were observed [Fig. S11] (see supplementary movie S2). In the first 50 s, two network bursts and nine subnetwork bursts appear. Seven of the nine subnetwork bursts were distinct burst patterns. Compared with the small-cluster network in Fig. 2, and the former two highly clustered networks in Fig. 4(a), where network bursts dominated the calcium dynamics, in this network, subnetwork patterns were more complex. Self-organized cultured neuronal networks rich in subnetwork burst patterns, though beneficial for illuminating the evolution from sporadic neuronal bursts to network bursts, were rarely documented [20]. In a well-synchronized network, random neuronal bursts can induce the synchronized burst of other neurons, hence explaining the dominant network bursts. However, in this network, only neurons 3 and 4 burst synchronously, indicating the formation of a strong bi-directional synaptic connection. No other neuron pair can promise a synchronized calcium activity. The failed neuronal burst transmission may indicate an unreliable information communication between the neurons [45].

It seems that networks with high aggregation degree can induce network bursts because of the more dense and reliable interneuronal synapses, while less clustered networks could potentially present richer subnetwork activity patterns. To examine this hypothesis, we resorted to patterned *μ*CP to confine neurons in desired areas and increase the aggregation degree. Meanwhile, other researchers have shown that symmetrically connected modular clustered networks also present dominant network bursts [20]. We wonder if asymmetry would break the reliable bidirectional signal transmission, hence allowing richer subnetwork bursts. To investigate this possibility at the same time, we changed the printing pattern to asymmetrically connected modular structures.

### C. Calcium dynamics in modular networks

Finally, we seeded primary neurons on agarose-treated PECM-patterned PS substrate, with the patterns being asymmetrically connected modular structures. In this group, the printed pattern contained two square modules connected by a belt of two parts: a cone-shaped part and a thin-line part [46]. This pattern was arrayed 10-by-12 to yield 120 identical patterns. Owing to the seeding randomness, we obtained both isolated networks (32/120) and asymmetrically connected modular networks (6/120) at one culture, all of which were highly clustered.

In the isolated networks, the hypothesis that stable network bursts could be induced by increasing the network’s aggregation degree was confirmed. Dominant network bursts reappeared in the highly clustered networks [Fig. 5] (see supplementary movie S3). More than that, other fascinating calcium behaviors were observed. First, contrary to previously reported observations [30, 37] that network bursts could not be induced in small-scale networks, a nine-neuron network in our culture presented dominant network bursts [Fig. 5(a2)]. Second, repeated network bursts were observed in a network as small as a two-neuron couple [Fig. 5(c)]. Another research by using micro-pillar substrate reported a similar two-neuron synchronized bursting behavior [47]. However, the information flow in that study was unidirectional, but not bidirectional. Moreover, an isolated neuron presented repeated spontaneous neuronal bursts. These repeated single-neuron bursts couldn’t be induced by the diffusion of neurotransmitters from the nearby modules, because it paced asynchronously from them (Fig. S12). These three neurons may be intrinsically bursting neurons, which are prevalent *in vivo* and can present a burst of spikes by a short stimulation or by noise like miniature excitatory postsynaptic potentials [41, 48, 49].

**FIG. 5.**
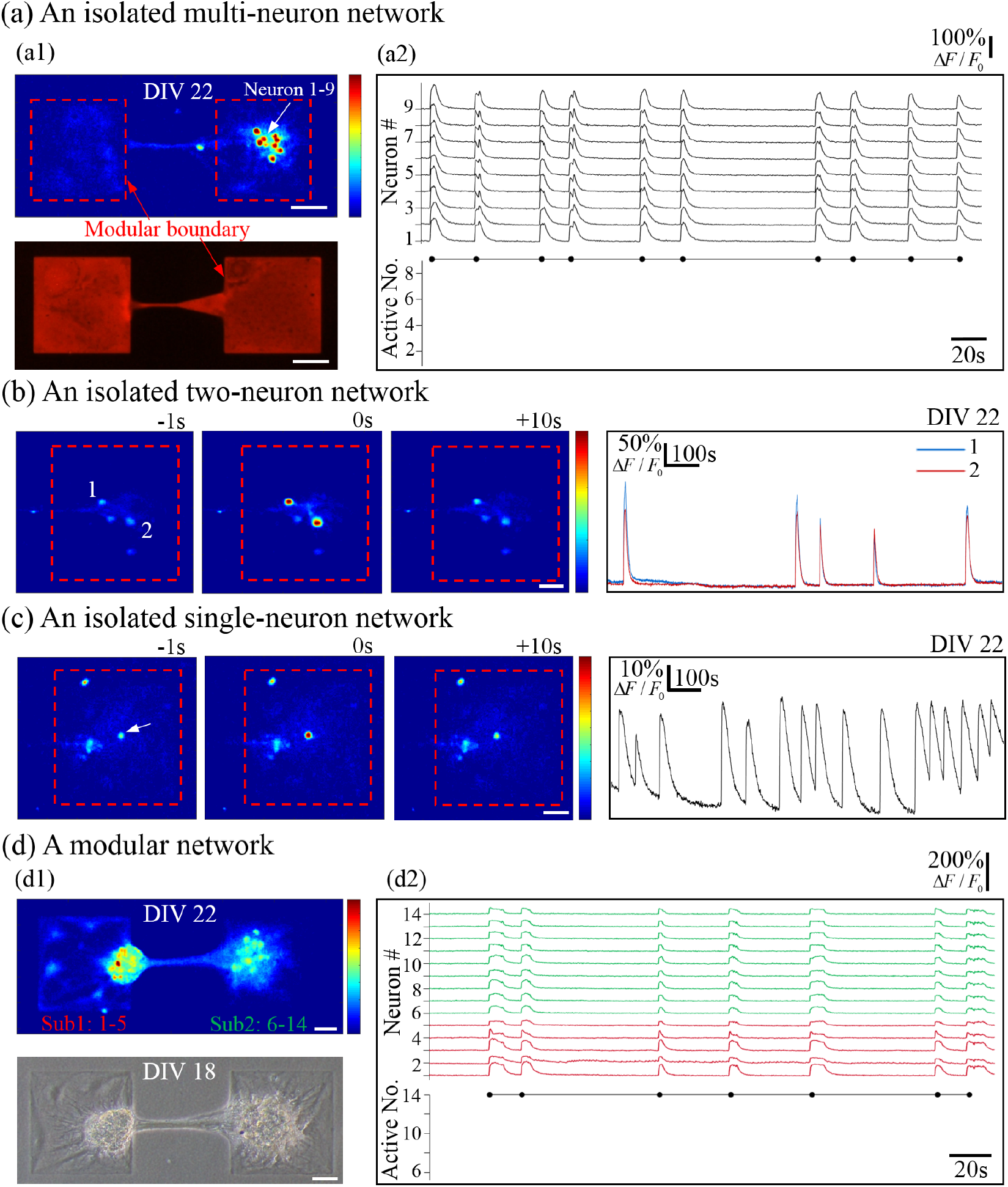
The locally-confined highly-clustered networks show dominant network bursts. (a1) False-color micrograph (top panel) of an isolated network, and its corresponding printed area (bottom panel), visualized by SR101. (a2) Top panel: relative fluorescence traces of neurons from (a1); bottom panel: active neuron number in a subnetwork burst versus time. (b) and (c) are from two isolated networks consisting of only two and one active neurons, respectively. Left three panels: False-color calcium recordings at a second before, right on, and ten seconds after a peak time, respectively; the rightmost panel: relative fluorescence traces of picked neurons (the loci indicated at the leftmost panels). (d1) False-color (top panel) and phase-contrast (bottom panel) micrographs of a asymmetrically-connected modular network. (d2) Top panel: color-coded traces of neurons in two subnetworks present synchronized neuronal bursts with post-burst plateau phases; bottom panel: active neuron number in a subnetwork burst versus time. Scale bar: 50 *μm*.

In the asymmetrically connected modular networks, two subnetworks were located on two separate modules, which were connected by a nerve bundle. Owing to the printed area’s limited neuron-substrate adhesion and high interneuronal attraction, neurons tended to reaggregate into small clusters within modules. Under the attraction force of neurites between two clusters, they were attracted to each other, even outside the square area [Fig. 5(d)]. However, in such an unusual circumstance, the network’s ability to transmit information was preserved, as evidenced by the repeated network bursts. Dominant network bursts were observed in all asymmetrically connected modular networks, as in symmetrically connected networks [20]. However, unlike in isolated, symmetrically connected, or small-cluster networks, where a short-term rapidly calcium-elevated phase was immediately followed by an exponentially decaying phase, in this asymmetrically connected modular network, an extended fluctuating plateau phase was inserted between the two phases. This could have been caused by the asymmetrical connection or the reverberatory signal transmission between modules [19]. Moreover, the duration of such a plateau phase between two consecutive network bursts greatly varies [Fig. 5(g)], indicating the variability of small-scale networks in calcium dynamics, consistent with other research [20, 37].

## III. CONCLUSION

In summary, we demonstrated that reduced cell-substrate adhesion caused by *μ*CP, higher seeding density, and local restriction would separately contribute to more aggregated networks, which were more likely to present dominant network bursts. On a uniform microcontact-printed substrate, densely seeded neurons formed a Petri dish-wide small-cluster network, of which the calcium dynamics was dominated by network bursts. After reducing the interneuronal attraction by diminishing the seeding density, the sparsely seeded neurons formed several networks with different clustering degrees. Less clustered networks presented time-varying subnetwork bursts. We then constructed highly clustered networks by confining neurons in small squares, where dominant network bursts reappeared. These findings demonstrate the existence of structure-dependent spontaneous calcium dynamics in cultured networks on microcontact-printed substrates, which provide insights into deciphering the initiation and propagation of synchronized network-wide bursting behaviors, which are prevalent in networks *in vitro*, *in silico*, and *in vivo*. Moreover, these findings also provide guidelines and implications for custom designing neuronal networks’ structure and functionality by using *μ*CP.

## IV. METHODS

### A. Processing of calcium fluorescence intensity recording videos

All recorded videos were processed preliminarily using ImageJ (National Institutes of Health) to produce the time courses of absolute fluorescence intensity of manually-selected regions of interest (ROIs). The recorded AVI files were imported and converted to grayscale image stacks to reduce the calculation complexity. Then, the brightness and contrast were adjusted to an appropriate level to make the boundary of neurons or clusters clear. After that, ROIs were selected as oval regions, and the fluorescence intensity traces were extracted, every point of which being the average intensity of a certain ROI on that frame.

The absolute fluorescence intensity traces of different ROIs were then fed into a custom Matlab script to output the relative intensity traces. First, the absolute fluorescence traces were filtered by a low-pass filter. Then, the intensity diminishing effect caused by photo-bleaching and fluorescence dye leaking was compensated by first fitting with an exponentially decaying function and then being subtracted by this function. To avoid the intensity fluctuation caused by neuronal bursts, the traces were sampled as the lowest points by a window of 50 s width at 50 s stride. These lowest points were then fitted by an exponentially decaying function with a non-linear least square method. Next, the background fluorescence intensity *F*_*i*,0_ was calculated as the median of the 20% lowest intensity points of each compensated trace. Finally, the relative fluorescence intensity traces of each ROI i was calculated as **f**_*i*_(*t*) = (**F**_*i*_ − *F*_*i*,0_)/*F*_*i*,0_, with *F_i_* being the absolute intensity trace of ROI *i*. Fig. S5 presents the calculated exemplary relative fluorescence intensity traces of different networks by the proposed method.

### B. Neuronal bursts detection and burst traces extraction

The onset and end time of each neuronal burst were inferred by a time-derivative thresholding method, which scanned the time derivative of the relative fluorescence traces. The onset time was labeled when the derivative reached +2.58 SD threshold from below, and the end time was labeled when the derivative reduced to zero from above after the onset time. The standard deviation (SD) for the derivative of each relative fluorescence trace was calculated after intercepting the derivative trace. The interception rule works as follows. First, the median of the derivative trace was determined. Then, a fixed-value range threshold (−0.1, +0.1) was offset by the median, and those not located in this range were eliminated. After that, SDs were calculated by the remaining derivative points. To reduce the false-positive detection results, the onset times of detected neuronal bursts were double-checked by a five-frame derivative. More than that, those detected neuronal bursts with no apparent relative fluorescence elevation were erased by using a fixed-value threshold (median + 0.1). The median here was calculated as the median of the lowest 20% points of the relative fluorescence traces, rather than the one mentioned in the last paragraph. Fig. S6 proved that the proposed neuronal burst detection method works well on the collected dataset.

### C. Subnetwork and network bursts detection

In all cases, subnetwork burst was inferred to exist when at least x neuronal bursts with onset time within a certain time window *t_win_* were detected. The number threshold *x* was 2 for all three groups of networks. The time window *t_win_* was 240 ms (3 frames) for small-cluster large-scale networks and weakly-separated medium-scale networks, but was 400 ms (5 frames) for strongly-separated small-scale networks. The choice of time window was optimized to avoid the undesirable division of apparent one subnetwork burst into two separate ones.

Considering the inevitable false-negative neuronal burst detection, any subnetwork burst in which at least 80% of all neurons in the network participate is inferred as a network-wide burst activity.

### D. Correlation coefficients

The correlation coefficient (CC) between neuron *i* and *j*, *r_ij_* was calculated using the respective relative fluorescence traces **f**_*i*_(*t*) and **f**_*j*_(*t*),

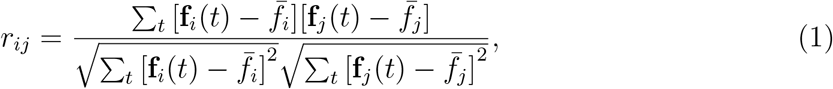

where 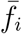 and 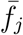 are the average relative intensity of the two traces, respectively. For each network or subnetwork, the mean CC within the network or subnetwork was computed as 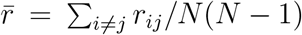, where *N* is the number of the network or subnetwork. The intra-subnetwork standard error of mean (SEM) was calculated as

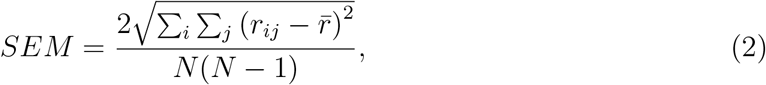

where *i < j*. The inter-subnetwork mean CC between two subnetworks was evaluated as, where *i* = 1,, *M*, and *j* = *M* + 1,, *N*. *M* and *N − M* are the number of the two subnetworks, respectively. The inter-subnetwork SEM was calculated as

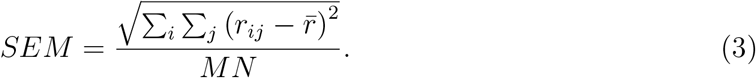

All statistical analyses were performed using the Student’s unpaired t-test. **Statistically significant difference with a p-value of < 0.01, “ns” indicates no statistical difference.

### E. Decay time constant

Every neuronal burst is followed by a decay phase, which represents the process of calcium ions being ejected from the cytoplasm. The speed of this process is quantified by the decay time constant, which was calculated by fitting the decay phase with an exponentially decaying function.

First, for each neuron, the decay phase after each neuronal burst was extracted and averaged [Fig. S7]. The overlapped decay phase of the relative fluorescence traces of a certain neuron was shown in Fig. S7(A), in which the end time of the last neuronal burst corresponds to the onset of the decay phase, and the onset time of the next neuronal burst represents the end of this decay phase. However, the inferred end time of a neuronal burst may not coincide with the onset time of the decay phase [Fig. S6(B)]. Considering that the decay phase usually starts from a peak, which may lie inside the burst or shortly after the end time of the burst, the onset time of each decay phase is inferred as such a peak time.The peak was located by a locally searching algorithm, which searched for the maximum in a time window Δ*t* followed by the end time of the last neuronal burst. The time window was locked to the next inter-burst interval *T_IBI_* by defining Δ*t* = *T_IBI_*/2 to balance computational efficiency and safety. The extracted traces were then averaged over time, with those time points containing only one trace being eliminated to reduce the noise.

Second, the averaged trace was fitted by an exponentially decaying function, which contains three unknown parameters: the initial state relative intensity *I_ini_*, the steady state intensity *I_ss_* and the decay time constant *τ*. The function is:

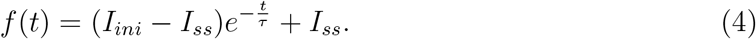

Third, the goodness of fit was calculated to evaluate the effect of fitting:

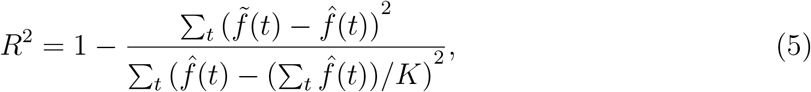

where 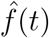 is the averaged trace, *K* is the number of the time points of the averaged trace, and 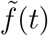 is the time-discretized form of the fitted function. The average and SEM of *R*^2^ were evaluated for the decay time constant of each neuron in the corresponding network. The fitting was also evaluated and visualized by comparing the fitted trace and the averaged trace in Fig. S7(B) and (D), where the good fitting effects were intuitively confirmed.

### F. Burst duration

Burst duration is defined as the duration of a certain neuronal burst, which was computed as the difference value between the onset and end time of the neuronal burst. The histograms of burst duration of all neuronal bursts were presented, as in Fig. S10. The bin width is the minimum time resolution, i.e., 80 ms. Left and right limits are 0−*bin*/2 and max(*duration*)+*bin*/2, respectively, where max(*duration*) represents the maximum duration of all neuronal bursts. The distribution of burst duration was fitted by a Gaussian function:

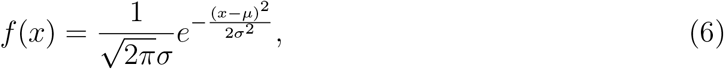

where *μ* and *σ* are two unknown parameters to be ascertained.

### G. Burst frequency

Burst frequency represents the average number of neuronal bursts in one neuron within one second, which was calculated as the number of all neuronal bursts of a certain time window, divided by the multiplication of the number of neurons and the duration of the time window. The width of the time window is set as 640 ms. The neuronal raster plot was scanned by the window at a stride of 80 ms to produce a time-dependent burst frequency vector. Average and SEM of burst frequency were calculated based on the vector to quantify the overall frequency and time-dependent fluctuation of burst activity, respectively.

### H. Inter-burst interval

IBI was computed as the difference value between the onset time of two consecutive neuronal bursts. The histograms of IBI of a certain experiment were presented, as in Fig. S8. All IBIs computed in an experiment were grouped into a vector *V_IBI_*. Then, the bin width of the histogram was set as [max(*V_IBI_*)−min(*V_IBI_*)]/*C*, where the constant *C* was optimized to produce the maximum *R*^2^ for Gaussian fitting considering the minimum temporal resolution, i.e., 80 ms. The left and right limits were 0 − *bin*/2 and max(*V_IBI_*) + *bin*/2, respectively. As in the burst duration analysis, the distribution of IBI was also fitted by a Gaussian function.

### I. Coefficient of variance of inter-burst interval

The coefficient of variance of IBI (IBICV) for a certain neuron was calculated as the standard deviation (SD) of IBI distribution divided by the mean of IBI. IBICV distribution can reveal the activity pattern of a network. For the noise-free deterministic or rhythmic activity, IBICV identically equals zero [Fig. S9 (A)]. For random Poisson activity, IBICV distributes normally with a mean value near 1 [Fig. S9 (B)]. In the simulations, 1000 independent neurons burst either rhythmically or randomly for 100 seconds at a mean burst frequency of 0.2 Hz. The time step was 0.1 ms. The histograms were presented and fitted by Gaussian functions, which revealed that either for simulations or for experiments, IBICV is distributed normally.

## V. ADDITIONAL INFORMATION

See supplementary material for details on materials and methods, for supplementary Figs. S1-S13, and for representative spontaneous calcium recordings of (1) small-cluster large-scale networks on unpatterned microcontact-printed substrates (Movie S1), (2) networks with different aggregation degrees induced by diminished interneuronal attraction (Movie S2), and (3) locally confined highly clustered networks (Movie S3).

## VI. ACKNOWLEDGEMENT

This work was supported by the National Natural Science Foundation of China (Grant Nos. 62022014 and 62173043).

## VII. DATA AVAILABILITY

The data that support the findings of this study are available from the corresponding author upon reasonable request.

## Supplementary Figures

**Fig. S1.**
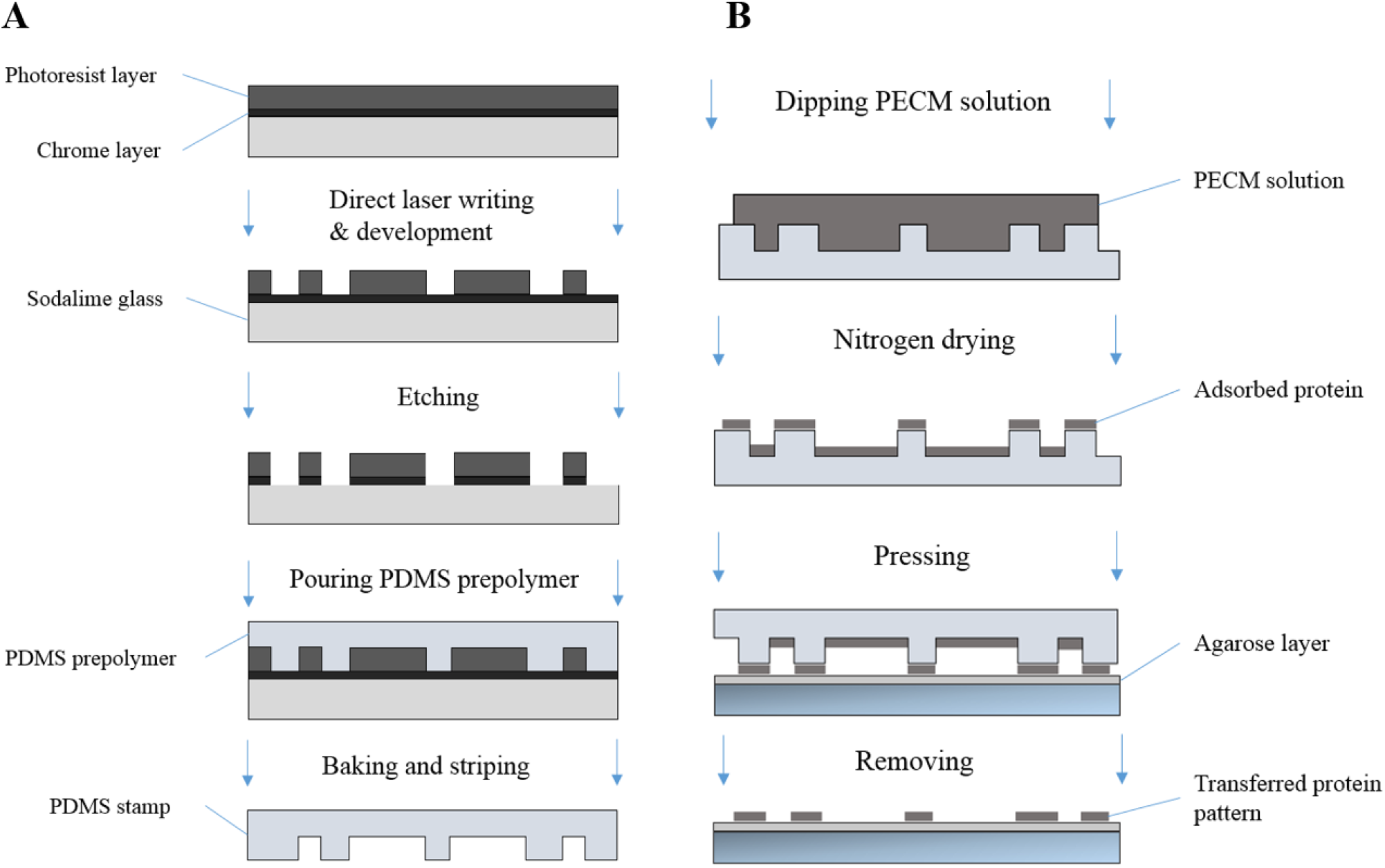
The processes of fabricating surface-modified protein-printed substrate. (**A**) Fabricating the PDMS micro-stamp. (**B**) Modifying and patterning the substrate.

**Fig. S2.**
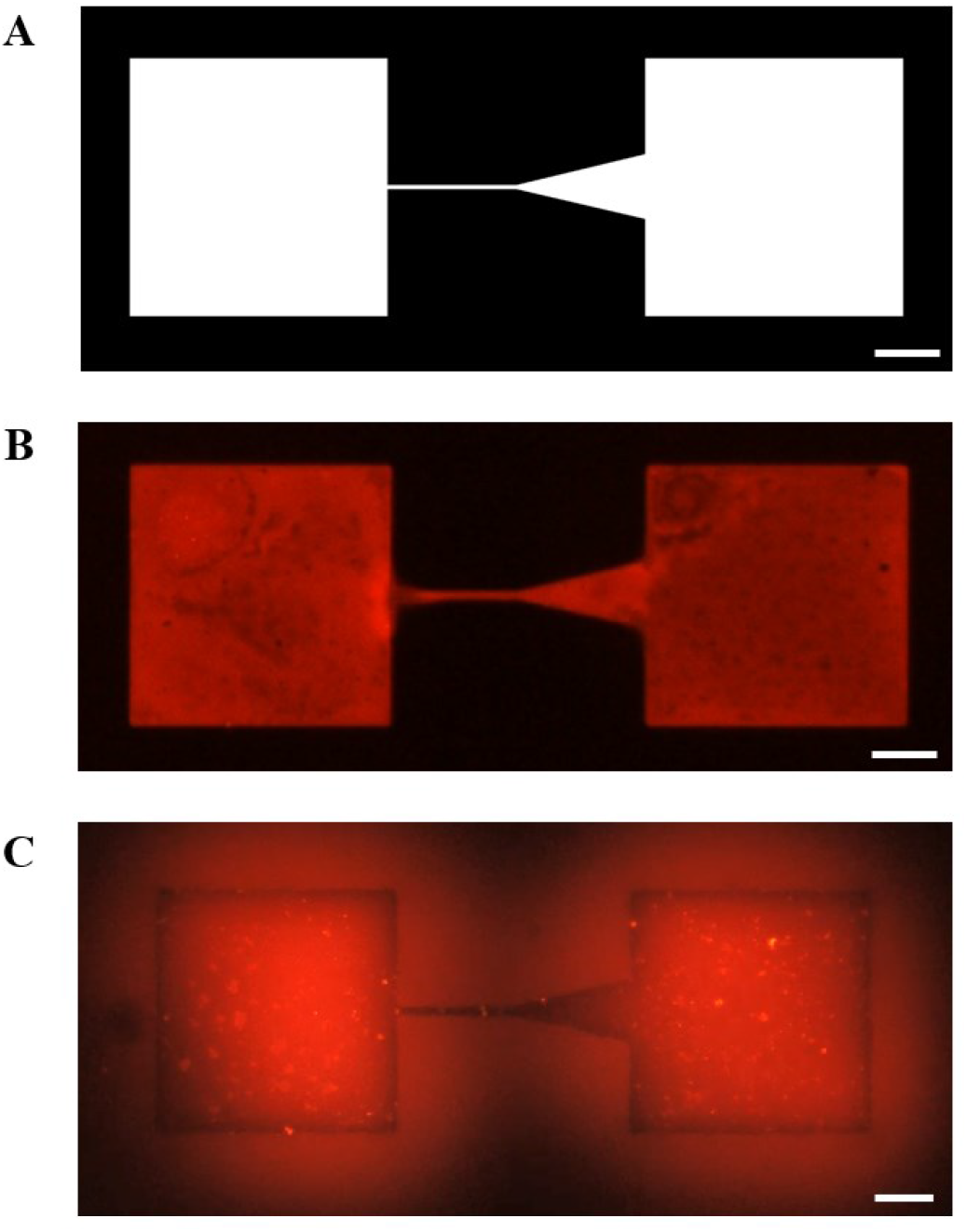
Evaluation of the pattern transfer effect by printing cell-permissive protein on the cell-nonpermissive modified substrate. (**A**) Digital image form of the designed pattern. (**B**) The fluorescence micrograph of the printed area presented a clear and desirable boundary, which confirmed the successful protein transfer by micro-contact printing. (**C**) Fluorescence micrograph of (**B**) after incubation at a humidified environment (relative humidity: 95%) for 1 hour. Under humidified environment, a thin layer of water film formed at the substrate’s surface. Adsorbed small-molecule fluorescence dyes were dissolved into the water film and diffused to surrounding unprinted area, while less soluble biomacromolecules remained attached to the printed pattern. Therefore, the printed protein area presented a black contour, which was surrounded by a red halo, which also demonstrated that the desired protein pattern was transferred. Scale bar: 50 μm.

**Fig. S3.**
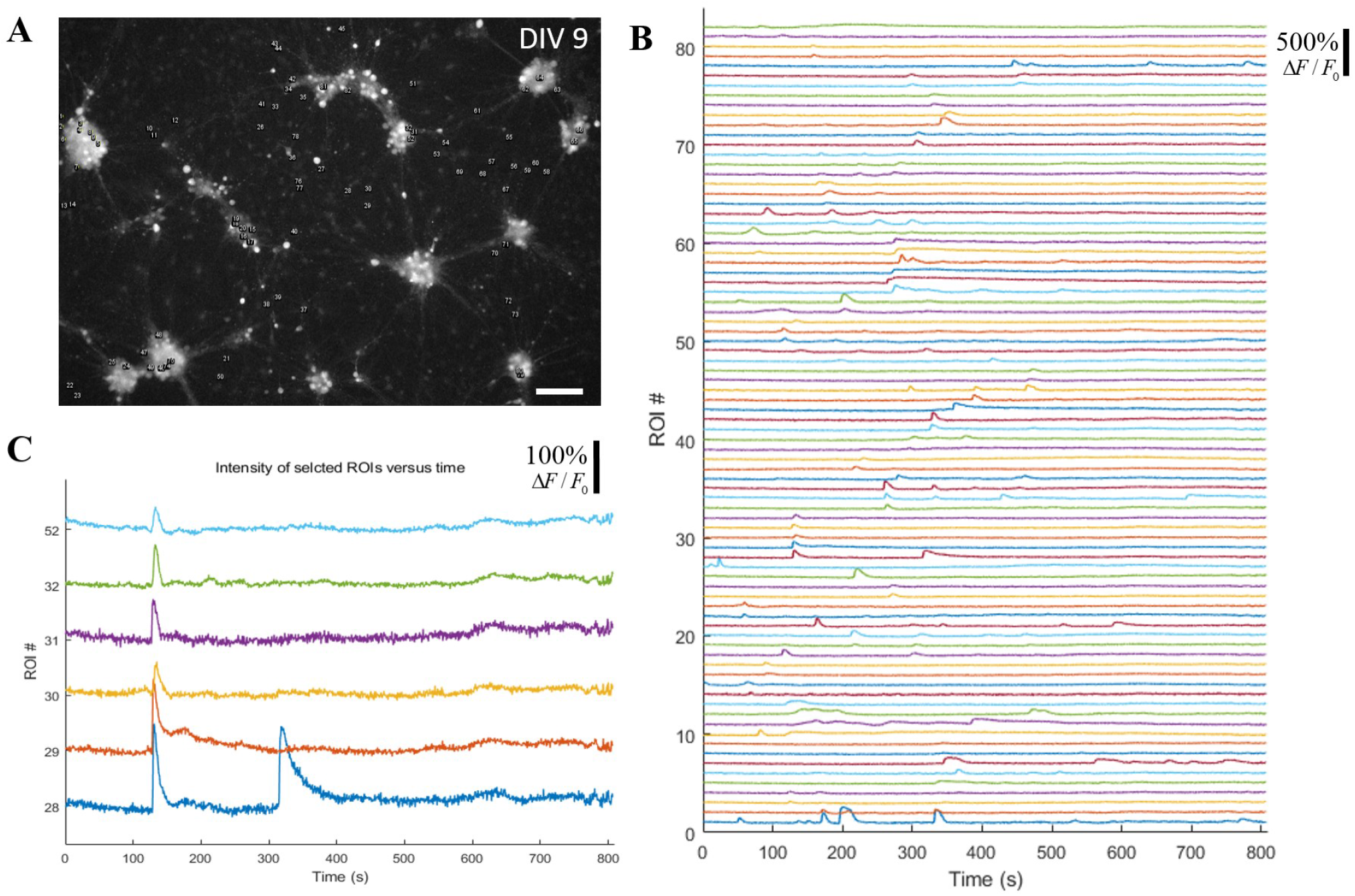
Only subnetwork-wide burst activities were observed in a small-cluster large-scale network on day 9. (**A**) ROI location in a small-cluster large-scale network on day 9. The grey image is the abstracted green channel of a peak-intensity frame of a calcium recording video. A total of 82 ROIs were selected manually. (**B**) Corresponding traces of relative fluorescence intensity of distinct ROI showed no network bursts, but only sporadic subnetwork bursts. (**C**) A six-neuron subnetwork synchronized calcium elevation on t = 140 s. Scale bar: 100 μm.

**Fig. S4.**
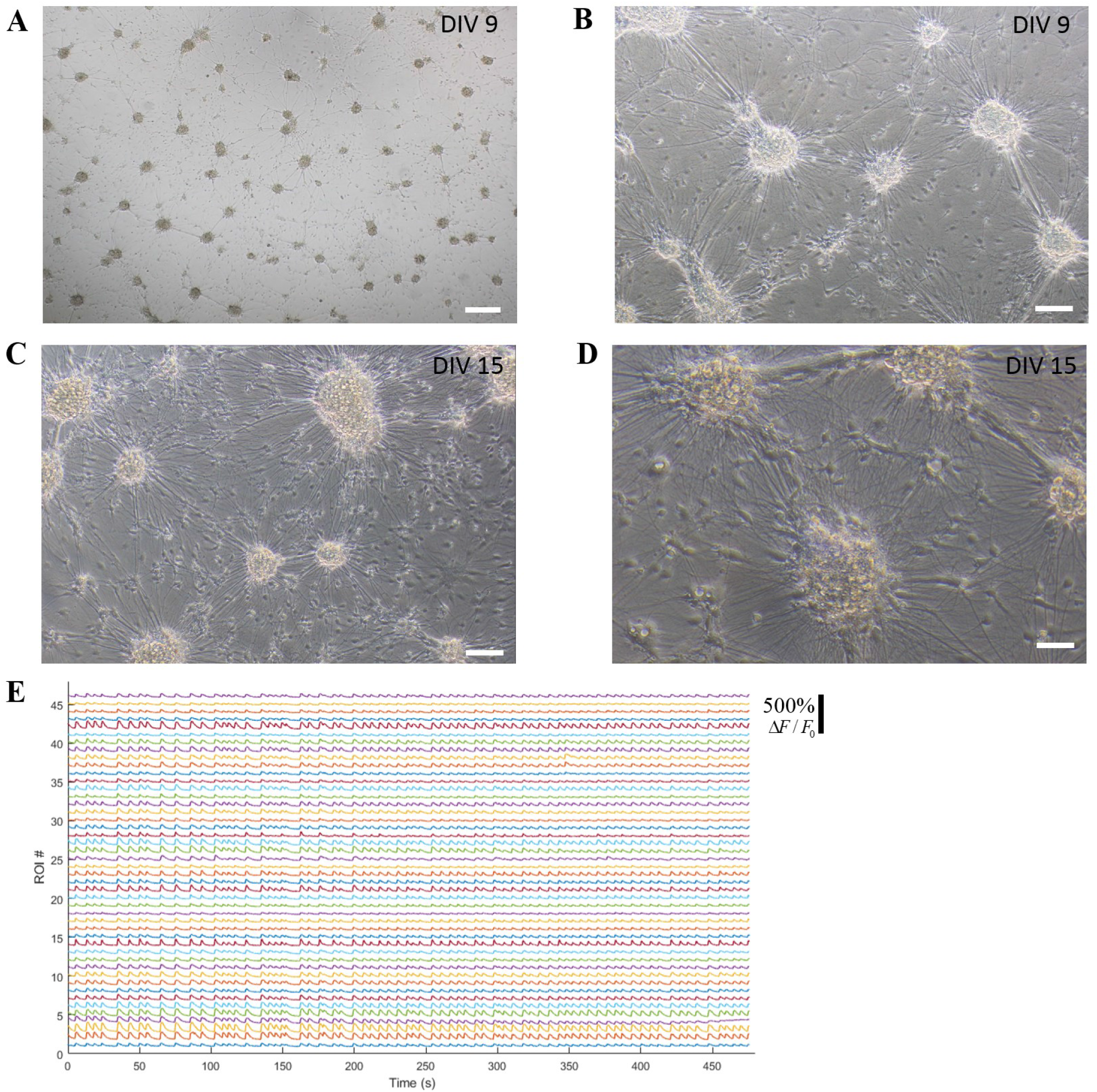
Network-wide burst activities and dense neurites were observed in a small-cluster large-scale network on day 15. (**A**) and (**B**) Phase-contrast micrographs of the small-cluster large-scale network on DIV 9. (**C**) and (**D**) Phase-contrast micrographs of the small-cluster network on DIV 15 revealed the formation of dense neurites. (E) Relative fluorescence of distinct ROIs in a small-cluster network on DIV 15 showed dominant network bursts. Scale bar: 250 μm, 100 μm, 100 μm and 50 μm in (**A**), (**B**), (**C**) and (**D**), respectively.

**Fig. S5.**
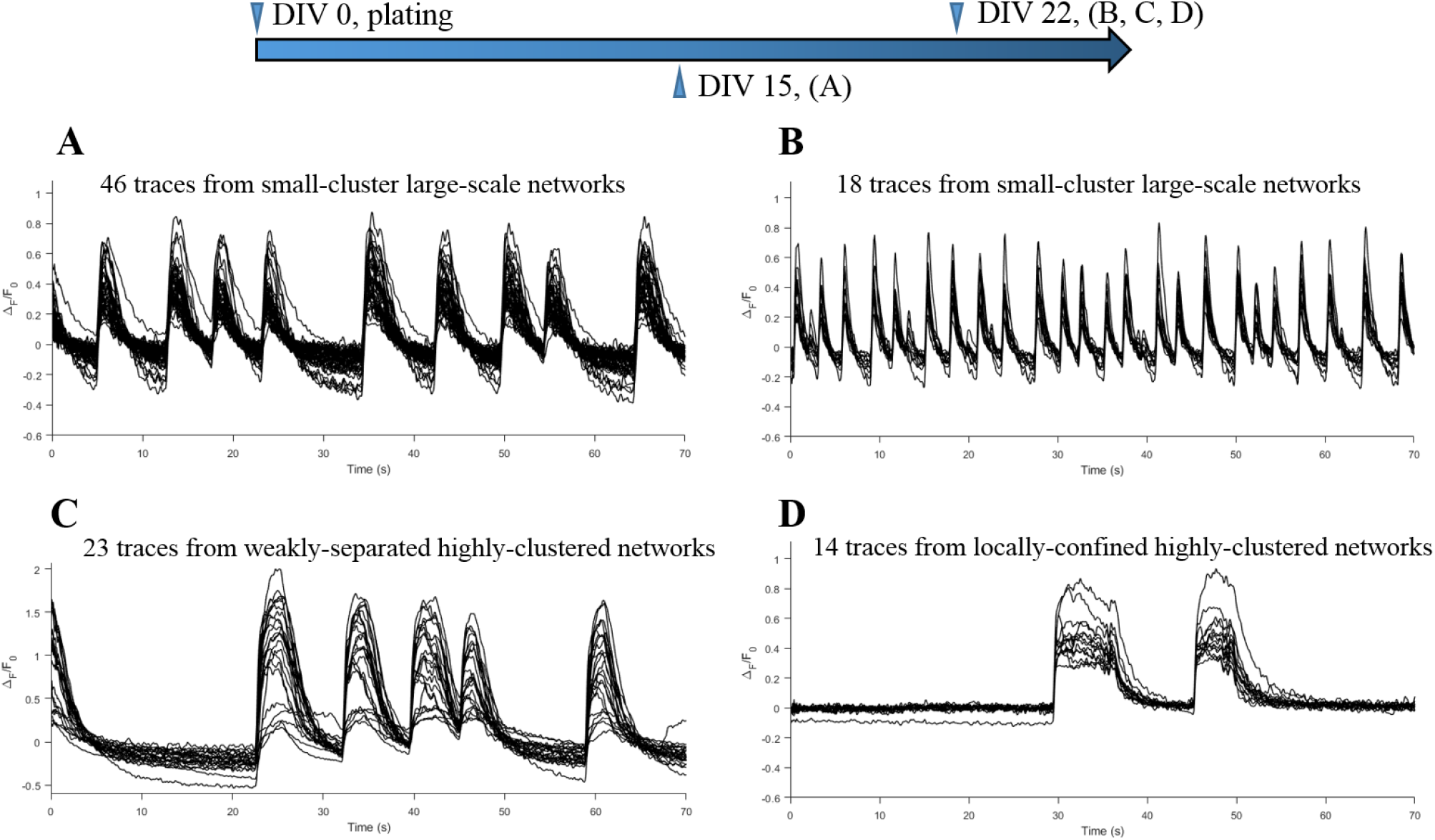
Representative overlapped relative fluorescence traces extracted from different cultured cortical networks. Traces from small-cluster large-scale networks on day 15 (**A**) and day 22 (**B**) show the high-synchronization of neuronal bursts within a network burst, corresponding to Fig. 2(a) and (b), respectively. Traces from selected synchronized neurons in a weakly-separated medium-scale network (**C**) and from a strongly-separated small-scale network (**D**) also show the synchronization of neuronal bursts within a network burst. (**C**) and (**D**) corresponds to Fig. 4(a) network 1 and Fig. 5(d), respectively.

**Fig. S6.**
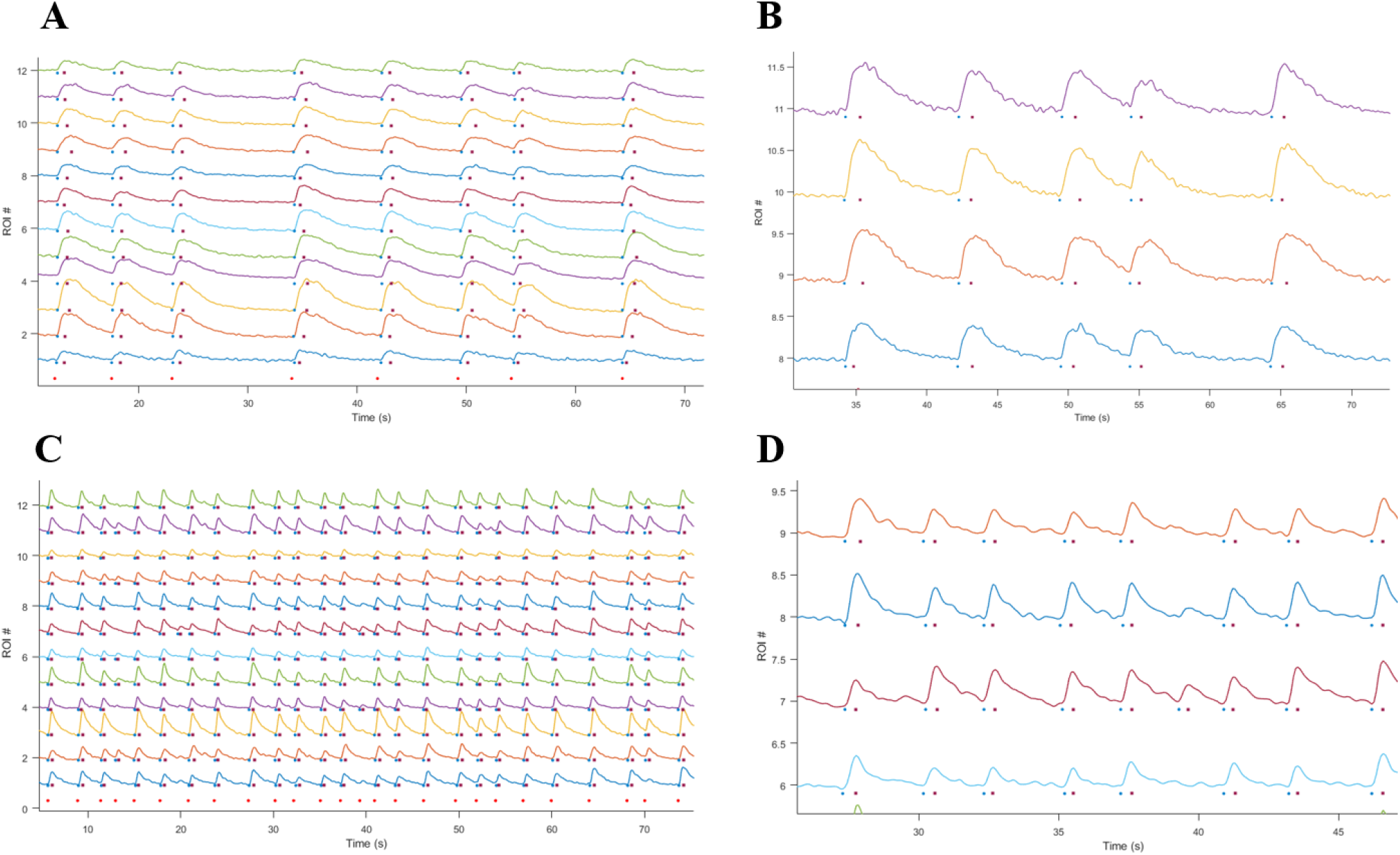
Illustration and evaluation of the burst extraction method. (**A**) Illustration of one example from a small-cluster large-scale network on day 15. (**B**) Zoom-in view of (**A**). (**C**) Illustration of another example from a small-cluster large-scale network on day 22. (**D**) Zoom-in view of (**C**). Blue round dots, dark-red square dots, and bright-red round dots represent the onset time of a neuronal burst, the end time of a neuronal burst, and the onset time of a subnetwork burst, respectively.

**Fig. S7.**
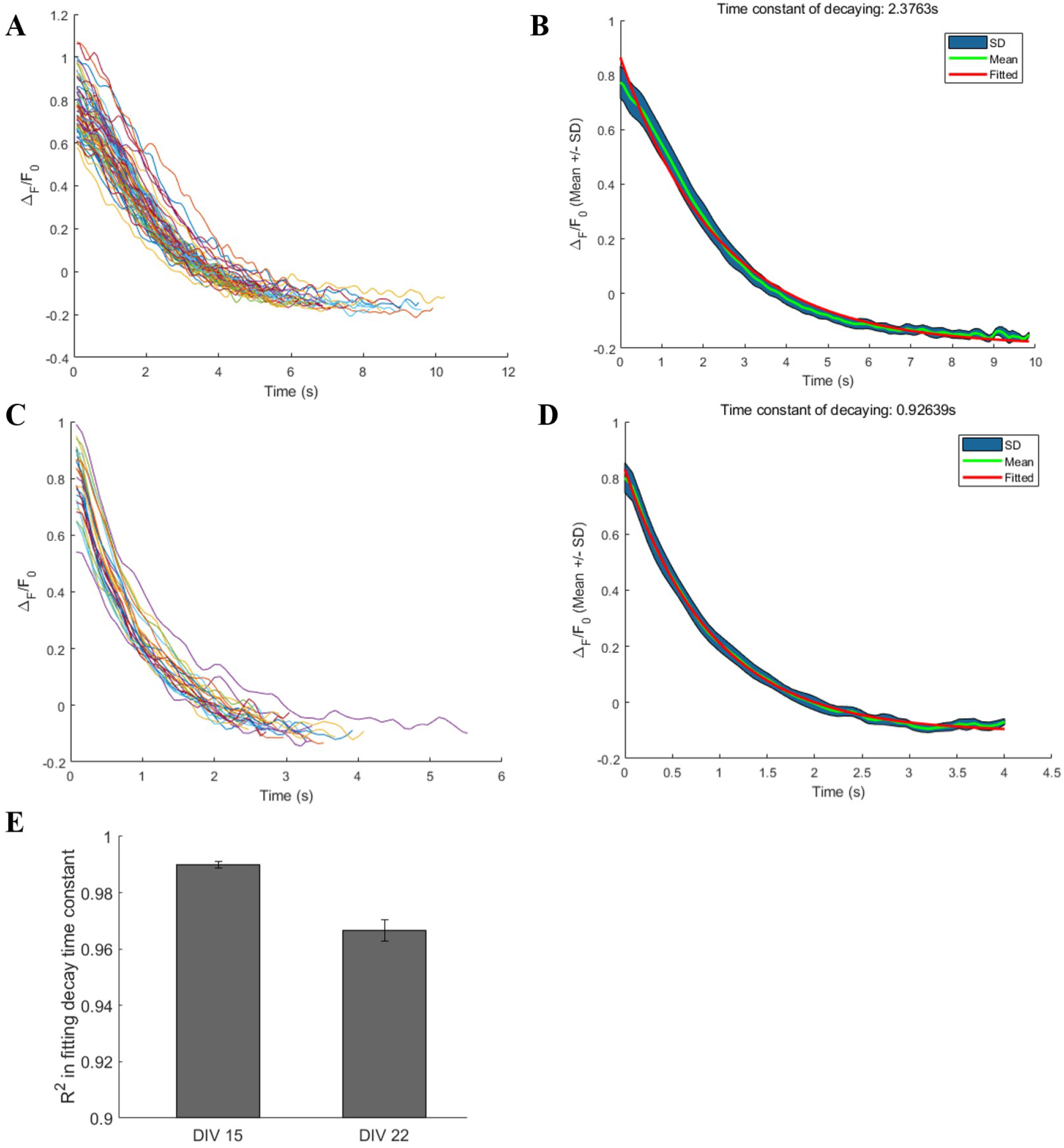
Extracting and fitting the calcium decay phase to calculate the decay time constant. (**A**) and (**C**) Extracted traces of two neurons from small-cluster large-scale networks on day 15 and day 22, respectively. (**B**) and (**D**) show corresponding traces of mean and standard deviation (SD) belt, traces of fitted exponential function, and the decay time constant. The goodness of fit R^2^ in (**B**) and (**D**) are 0.9909, and 0.9863, respectively. (**E**) R^2^ in fitting the time constant of decaying phase close to 1 on both day 15 and day 22 demonstrated the feasibility of the choice of exponentially decaying function and the good performance of the proposed method for extracting the decay phase. Data are presented with mean ± SEM.

**Fig. S8.**
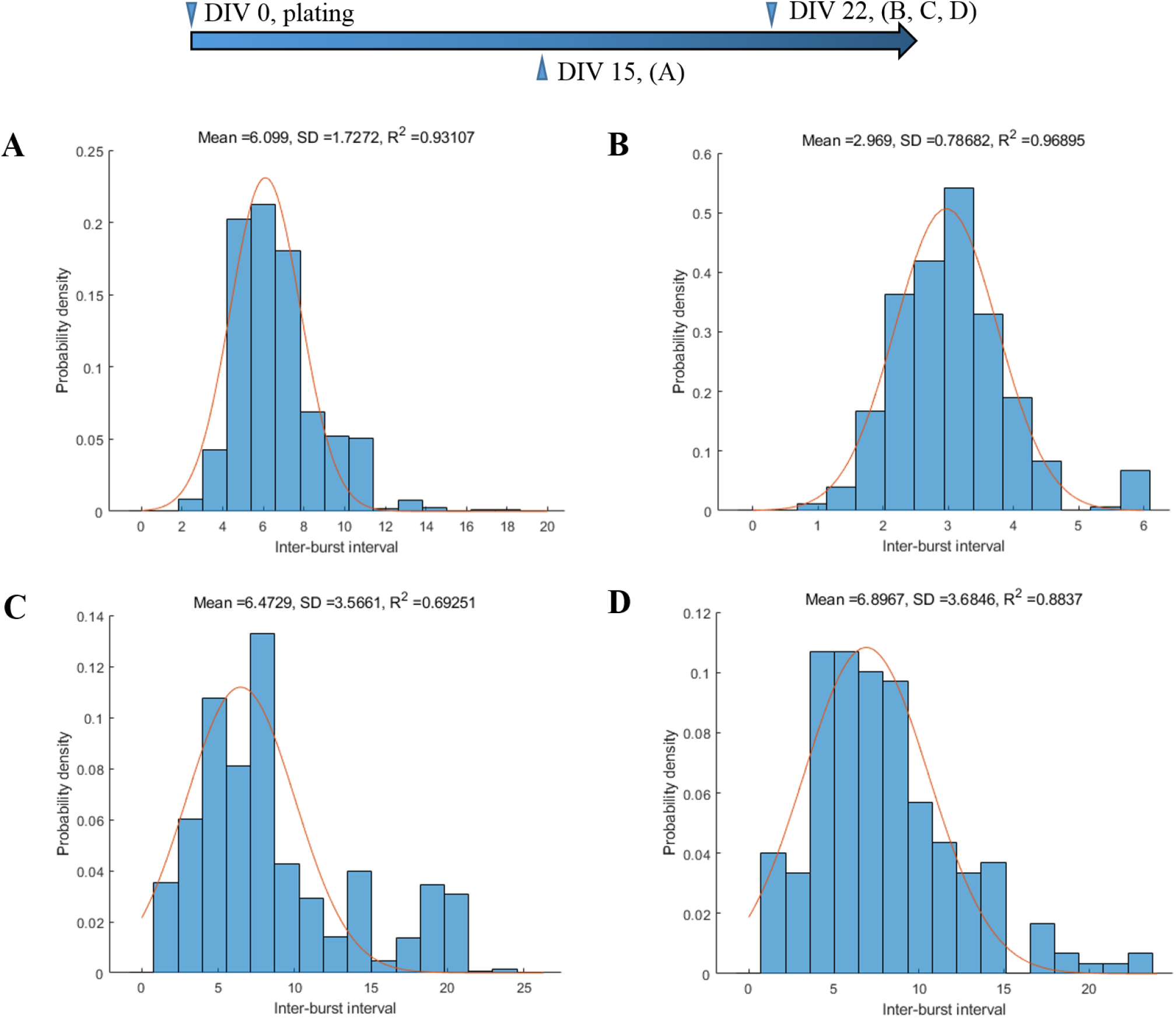
The distribution of inter-burst interval (IBI) of different cultured cortical networks. (**A**) A small-cluster large-scale network on day 15. (**B**) A small-cluster large-scale network on day 22. (**C**) Two weakly-separated highly-clustered networks taken as one merged network on day 22, as in Fig 4(a). (**D**) Another weakly-separated less-clustered network on day 22, as in Fig 4(b). The high values of *R*^2^ in (**A**) and (**B**) revealed that the IBI of the small-cluster large-scale network distribute normally. The relatively low value of *R*^2^ in (**C**) indicated that the IBI in the merged network distribute non-normally, probably due to the fact that the two subnetworks burst asynchronously. However, in the weakly-separated less-clustered network presenting abundant subnetwork activities, the IBI distribute near normally. The orange curves represent the fitted Gaussian function, with the fitted mean, SD and *R*^2^ presented above each subfigure.

**Fig. S9.**
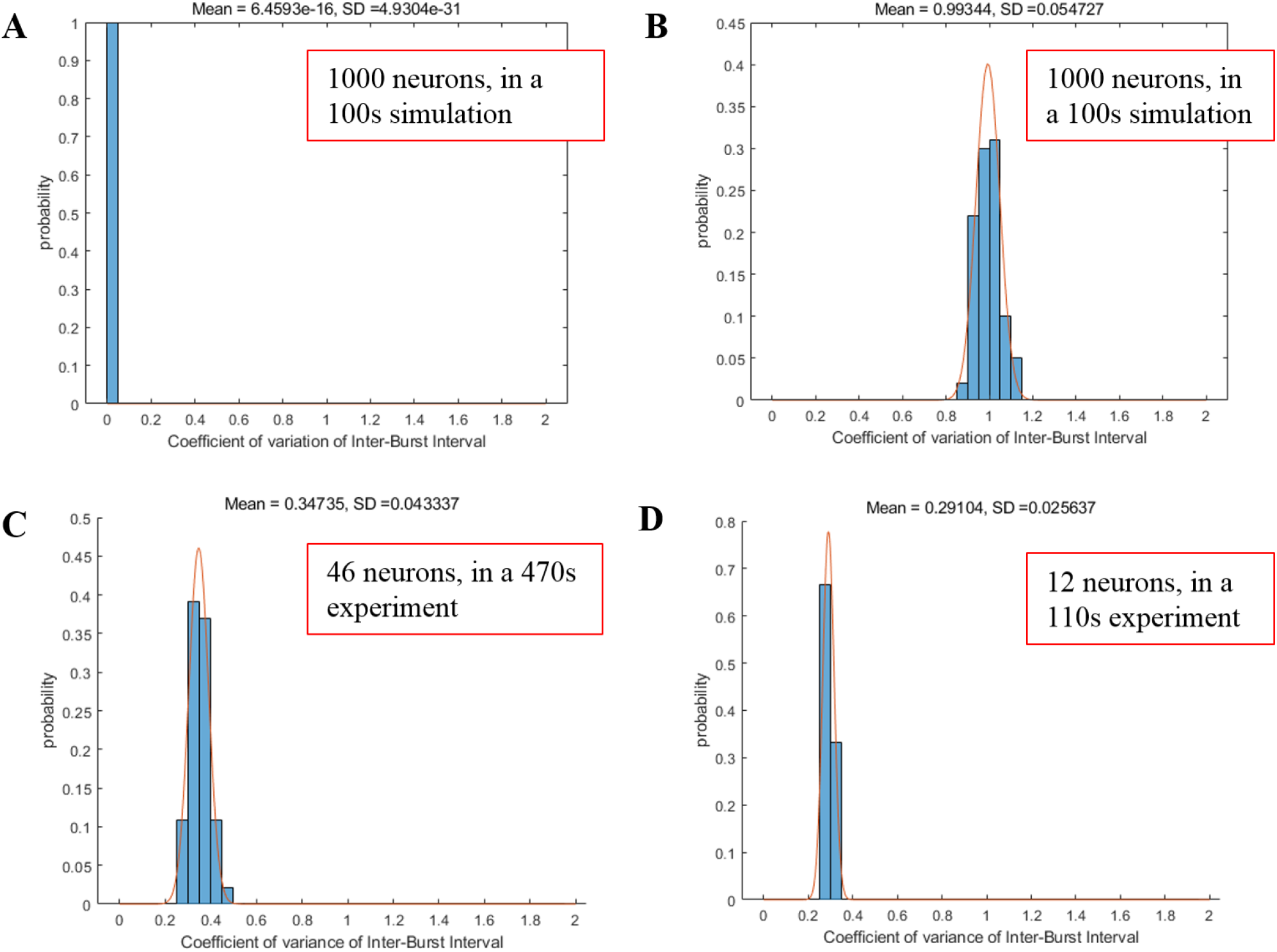
The coefficient of variance (CV) of IBI of different cultured cortical networks. (**A**) IBICV distribution of a deterministic process in a simulation consists of 1000 neurons with a duration of 100 s. (**B**) IBICV distribution of a Poisson process in a simulation consists of 1000 neurons with a duration of 100 s. (**C**) IBICV distribution of a small-cluster large-scale network on day 15, consisting of 46 recorded neurons with a recording duration of 470 s. (**D**) IBICV distribution of a small-cluster large-scale network on day 22, consisting of 12 recorded neurons with a duration of 110 s. The results revealed that small-cluster networks burst with a combination of a deterministic (rhythmic) process and a Poisson process. The orange curves represent the fitted Gaussian function, with the fitted mean and SD presented above each subfigure.

**Fig. S10.**
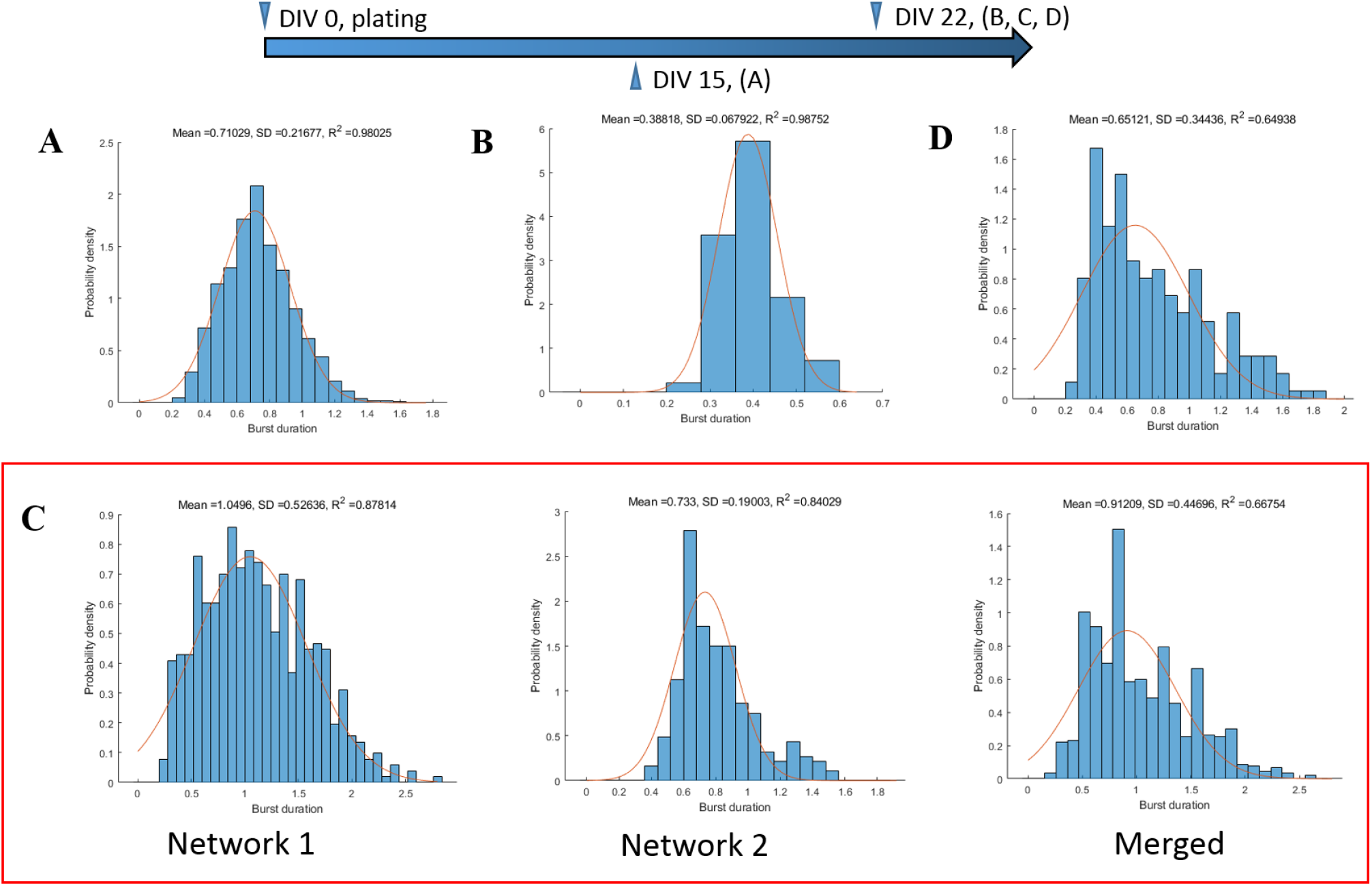
The distribution of burst duration of different cultured cortical networks. (**A**) A small-cluster large-scale network on day 15. (**B**) A small-cluster network on day 22. (**C**, left) Network 1, (**C**, middle) network 2, and (**C**, right) the merged one from two neighboring weakly separated networks with high clustering degree on day 22, as in Fig 4(a). (**D**) Another weakly separated network with low clustering degree on day 22, as in Fig. 4 (b). The high values of *R*^2^ in (**A**) and (**B**) showed that the burst duration in the small-cluster network was distributed normally. The difference of *R*^2^ in (**C**) indicated the burst duration of the two networks distributed normally but differently from each other. The low value of *R*^2^ in (**D**) suggested that the burst duration of this network was distributed non-normally. The orange curves represent the fitted Gaussian function, with the fitted mean, SD and *R*^2^ presented above each subfigure.

**Fig. S11.**
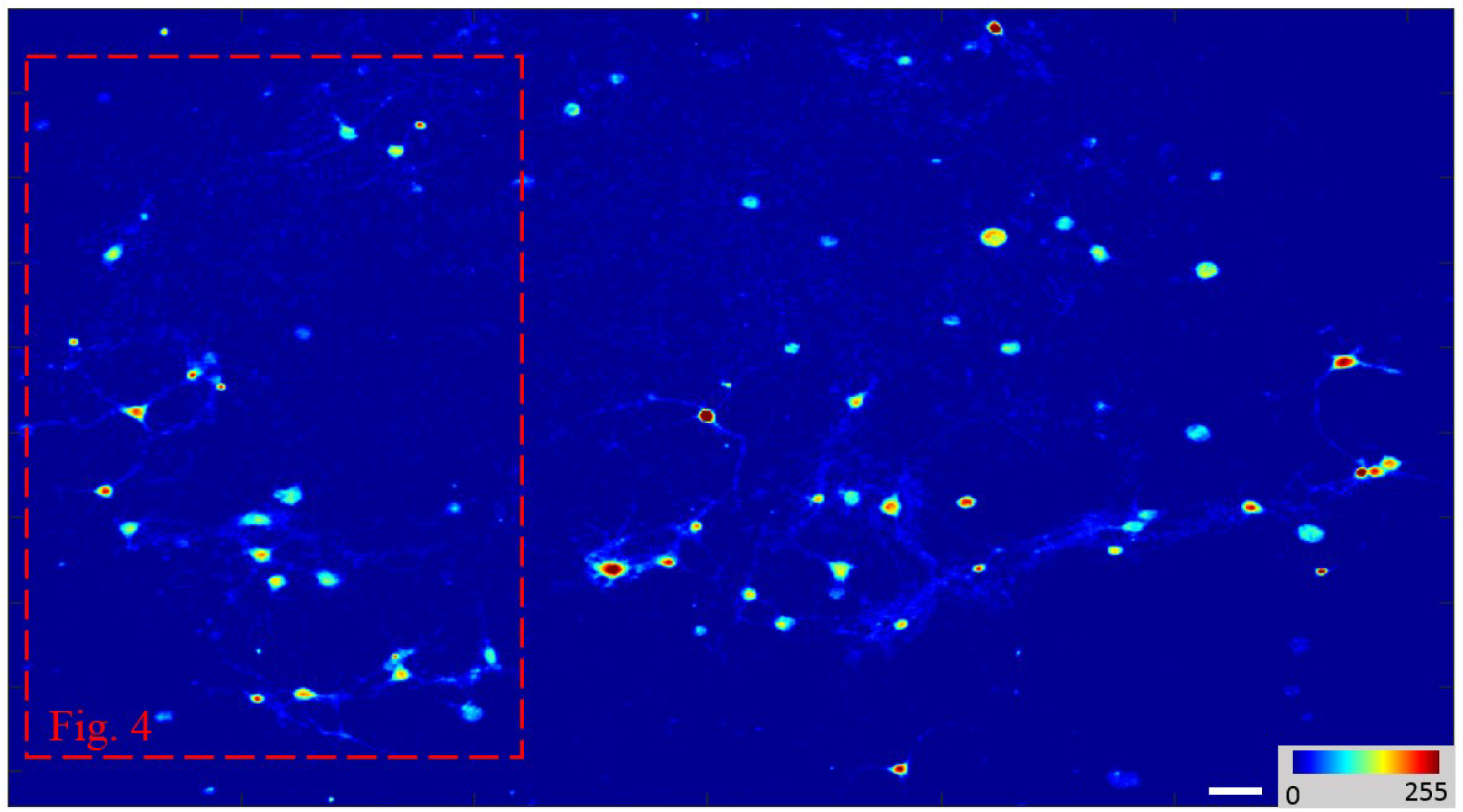
Fig. 4 is a subnetwork abstracted from a weakly separated less clustered network. Scale bar: 40 μm.

**Fig. S12.**
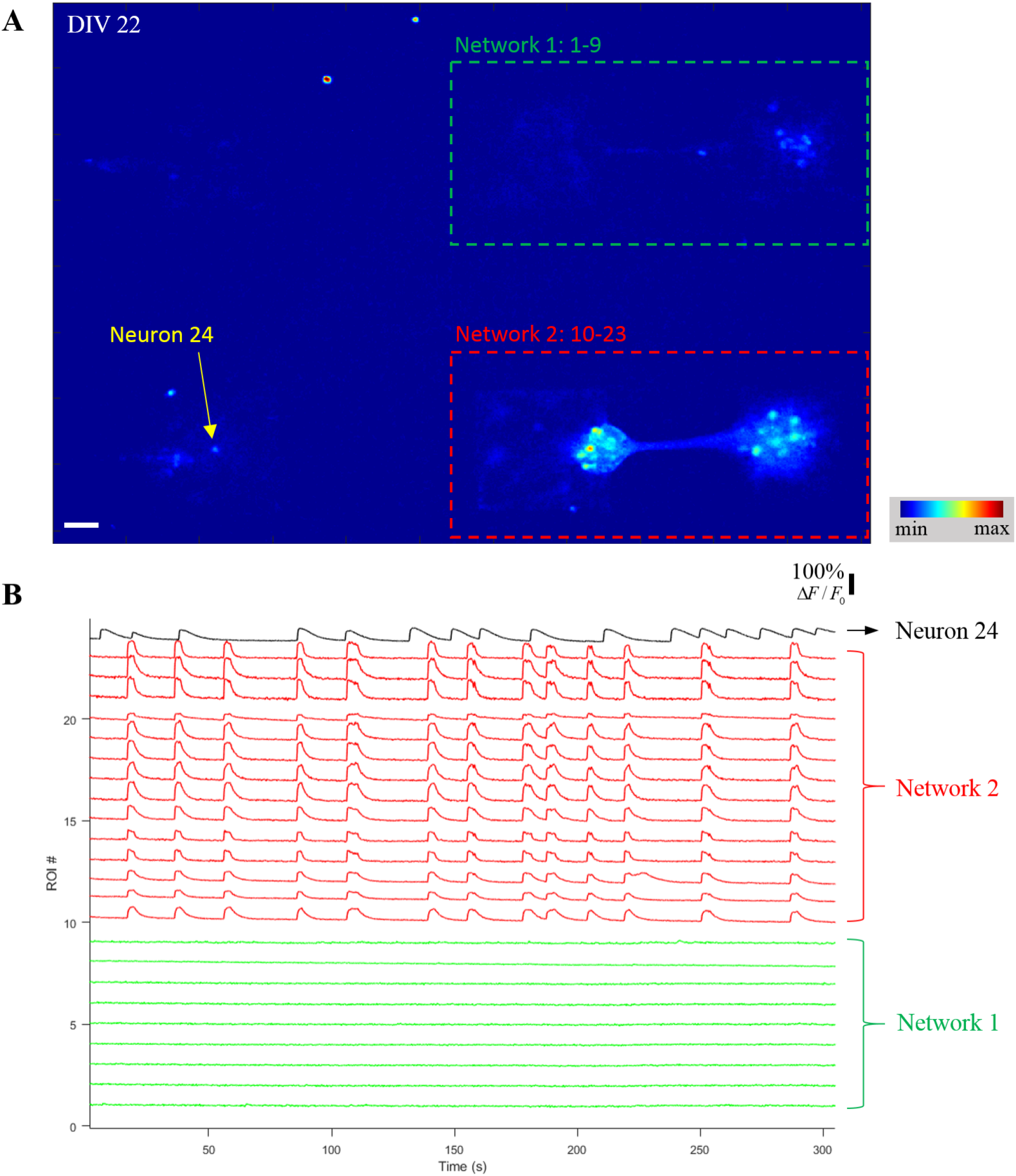
An isolated and spontaneously bursting neuron bursts asynchronously with two nearby separated neuronal networks. (**A**) False-color fluorescence micrograph of three strongly-separated small-scale networks. (**B**) Corresponding relative-intensity traces on different ROIs from (**A**). The isolated and spontaneously bursting neuron (ROI 24) was labeled with a black trace. Scale bar: 50μm.

## Notes

### Competing Interest Statement

The authors have declared no competing interest.

### Summary of Updates

Titles of reference articles added; Supplementary files updated.

