## Supplementary Notes and Figures S1-S12 for "Structure-dependent spontaneous calcium dynamics in cultured neuronal networks on microcontact-printed substrates"

**This PDF file includes:**

Supplementary Notes
Figs. S1 to S12
Legend for movies S1 to S3
References

Other Supplementary Material for this manuscript includes the following:

Movies S1 to S3

### Supplementary Notes

#### Fabrication of PDMS micro-stamps

Stamps with two kinds of patterns were fabricated, including unpatterned stamps (with a uniform surface) and patterned stamps (with raised 2-by-1 modular micro-patterns). First, chrome plates with a magnetron-sputtered chrome layer of 100 nm thickness and a pre-coated positive photoresist AZ4562 (AZ Electronic Materials Inc.) of thickness 12 μm on soda-lime glass were purchased from Shaoguang Inc. Second, the chrome plate was cut into small square pieces of 25 mm width by a glazier's diamond. Third, the small chrome piece was put and fixed on the vacuum holder platform of a Pattern Generator (microPG 101 from Heidelberg Inc.). Desired patterns were loaded and the direct laser writing process was initiated. Fourth, the exposed area of the photoresist layer was removed by developer AZ400K (AZ Electronic Materials) 1:3 mixed with ultra-pure water for 8 min, and then the exposed area of the chrome layer was also removed by Chrome Etchant 18 (Dow Materials Science Inc.) for 1min. Each step was followed by rinsing with ultra-pure water three times. Fifth, a polydimethylsiloxane (PDMS) stamp was fabricated by replica moulding. Before the developed and etched chrome plate was put in a 35 mm Petri dish, two layers of aluminum foil were put on a 35 mm Petri dish, attaching closely to the bottom and wall, to facilitate the separation process. PDMS pre-polymer (Sylgard 184 from Dow Corning Inc.) 10:1 mixed with curing agent was degassed at 5℃ for 30 min, and poured onto the Petri dish, followed by baking at 65℃ for 2 hours. Finally, the cured PDMS stamp was released from the chrome plate and cut into the desired shape. The fabricated PDMS stamps were stored in ultra-pure water when not used to increase the hydrophilicity.

#### Fabrication of surface-modified protein-printed substrates

First, the substrate of a 35 mm Tissue Culture-treated Petri dish (Dow Corning Inc.) was pre-coated by 0.2% agarose (BioFroxx Inc.) solution (in ultra-pure water) to render the substrate cell-repellent. The pipetted solution should avoid touching the dish wall to render a relatively uniform thin layer of agarose gel. Second, protein ink [1% ECM gel (E1270 from Sigma-Aldrich Inc.) + 50 μg/mL poly-D-lysine (PDL, P0899 from Sigma-Aldrich Inc.)] (called PECM) in PBS buffer solution (Thermo Fisher Scientific Inc.)] was prepared. The whole process should be guaranteed to be done aseptically. Filtering through a 220 nm filter is not allowed, because this would significantly reduce the protein concentration. Third, the stamp was pre-treated by coating a layer of sodium dodecyl sulfate (SDS, from Solarbio Inc.) to facilitate the protein transfer process. Before use, the PDMS stamp was put into a 10% SDS modification solution (in ultra-pure water). The solution was then sonicated for 5min and put standing for 5 min [1]. After that, the stamp was taken out of modification solution, dried by nitrogen, dipped into ultra-pure water to remove excess SDS, and dried by nitrogen again. Fourth, the inking process. The modified stamp was put inverted on a clean Petri dish. 200 μL PECM solution was pipetted onto the surface to cover the raised pattern area. The dish was covered and put on a 37℃

incubator for 20 min. Then, the stamp was taken out and dried by nitrogen quickly to avoid the formation of salt crystallization, which would greatly deteriorate the subsequent protein transfer process. Finally, the printing process. The agarose-coated substrate was sterilized by ultra-violet radiation for an hour. Then, the stamp was once again inverted and pressed against the agarose-coated substrate, with raised pattern against the substrate. A pressure of 300 g was applied for 2 min. After that, the stamp was released and the agarose-coated protein-printed substrate was obtained. The modified Petri dish was stored in a sterile clean bench. On the day before use, the Petri dish was added with primary neuron culture solution overnight and observed the next day to rule out the possibility of bacterial infection.

**Visualization of printed protein pattern**

10μg/mL red fluorescence dye Sulforhodamine 101 (SR 101, S7635 from Sigma-Aldrich Inc.) was added to the PECM solution to aid the visualization of the protein transfer effect. In Fig. S2(B), a fluorescence micrograph after printing revealed that the protein transfer process was desirable. After that, the printed Petri dish was put into a humidified incubator with 95% relative humidity for an hour, and examined again under a fluorescence microscope [Fig. S2(C)].

**Cell culture**

All experiments were approved by the Institutional Animal Care and Use Committee of the Beijing Institute of Technology. Primary cortical neurons (OriCell SCCFN-00001 from Cyagen Inc.) from E18 Sprague Dawley rats were dissociated, cryopreserved in preservation medium (Neurobasal medium (Gibco) + 10% DMSO), transported on dry ice, and preserved in liquid nitrogen before use.

Right before plating, the cryopreserved primary neurons in 1 mL preservation medium were quickly thawed and added with 4 mL pre-warmed neuronal plating medium (MEM + 5% fetal bovine serum + 5% horse serum + 0.6% D-glucose). Then, neurons were plated at desired density on agarose-modified PECM-printed substrates. The neuronal seeding of high-density and low-density mentioned in the main text are 600 and 150 cells $mm^{-2}$, respectively. After 4 hours, 2/3 medium was changed by neuronal culture medium (Neurobasal medium (Gibco) + 2% B27 + 1% GlutaMAX). Half the medium was changed every 4days since day 4 with neuronal culture medium. No anti-mitosis drug was added to inhibit the proliferation of neuroglia cells because they are essential to the survival of the barely anchored neurons and many neurons started to extend neurites only after the formation of glia carpet.

**Calcium imaging**

Calcium imaging experiments were conducted on day 9, day 15, and day 22. In each experiment, cultured neurons were loaded with 4 μM calcium indicator Fluo-4 AM (Solarbio Inc.) and 0.01% Pluronic F-127 in Mg2+-free buffered saline solution (BSS, containing 130 mM NaCl, 5.4 mM KCl,

5.5 mM Glucose, 20 mM HEPES, and 1.8 mM $CaCl_2$) at 37℃ for 40min, rinsed by BSS, and incubated again in BSS for another 10min before observation [2]. The cultures loaded with calcium indicator were observed on an inverted microscope (IX73, Olympus) equipped with a 10x objective lens, a short-arc lamp (X-Cite® 120Q, Lumen Dynamics Inc.), and a charge coupled device camera (DP21, Olympus). All calcium imaging videos were recorded by the cellSens software (Olympus) at room temperature, with a temporal resolution of 80 ms (12.5 frames/s) and spatial resolution of 1200x800 $pixel^2$. The faintest level of excitation fluorescence was selected to reduce the adverse effects of photo-bleaching and photo-toxicity.

For the small-cluster large-scale network, the same network was imaged at two different time points (on day 15 and day 22). For other experiments, there were no repetitive usages on different days.

### Supplementary Figures

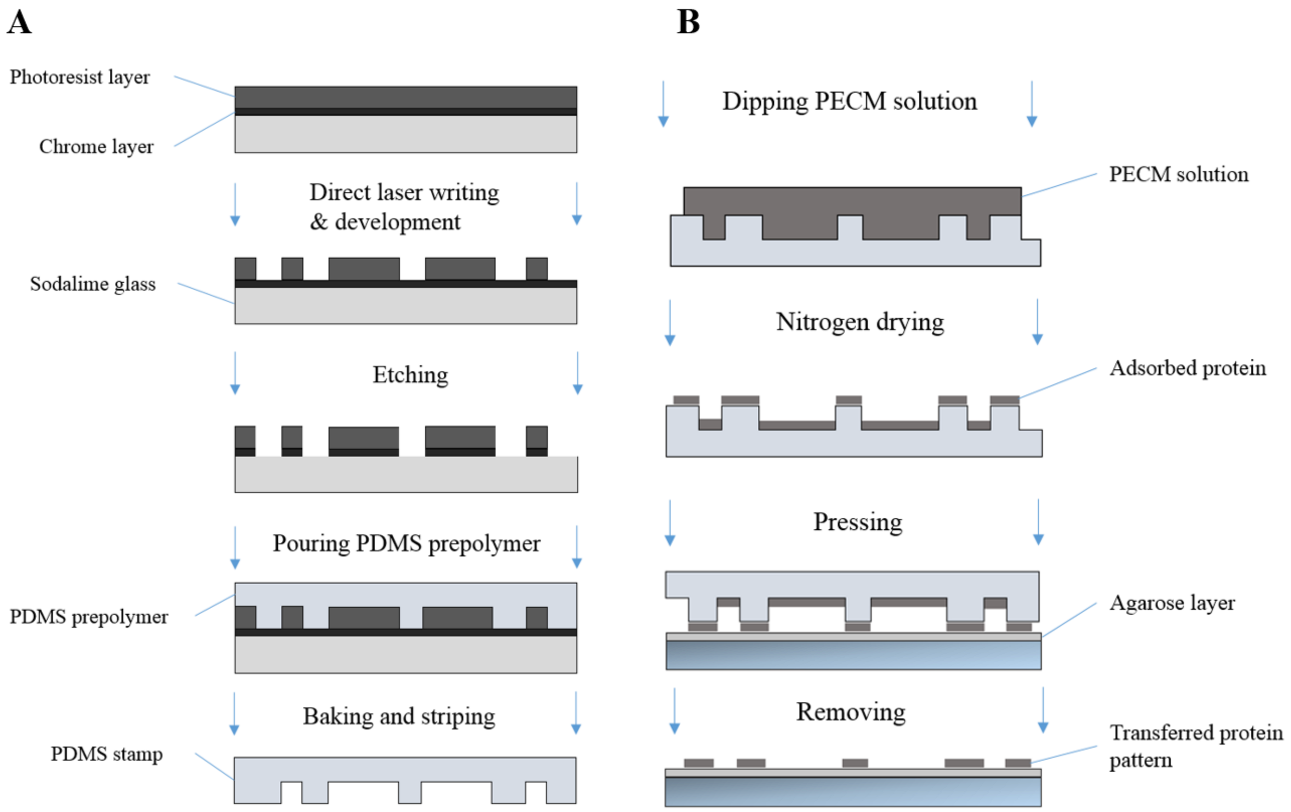


**Fig. S1. The processes of fabricating surface-modified protein-printed substrate.** (**A**) Fabricating the PDMS micro-stamp. (**B**) Modifying and patterning the substrate.

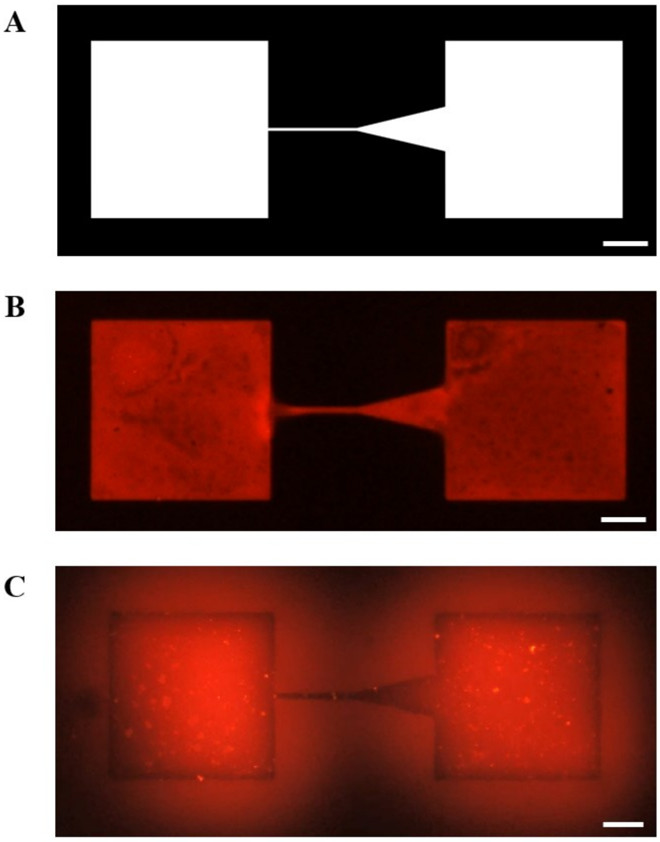


**Fig. S2. Evaluation of the pattern transfer effect by printing cell-permissive protein on the cell-nonpermissive modified substrate.**

(**A**) Digital image form of the designed pattern. (**B**) The fluorescence micrograph of the printed area presented a clear and desirable boundary, which confirmed the successful protein transfer by micro-contact printing. (**C**) Fluorescence micrograph of (**B**) after incubation at a humidified environment (relative humidity: 95%) for 1 hour. Under humidified environment, a thin layer of water film formed at the substrate's surface. Adsorbed small-molecule fluorescence dyes were dissolved into the water film and diffused to surrounding unprinted area, while less soluble biomacromolecules remained attached to the printed pattern. Therefore, the printed protein area presented a black contour, which was surrounded by a red halo, which also demonstrated that the desired protein pattern was transferred. Scale bar: 50 μm.

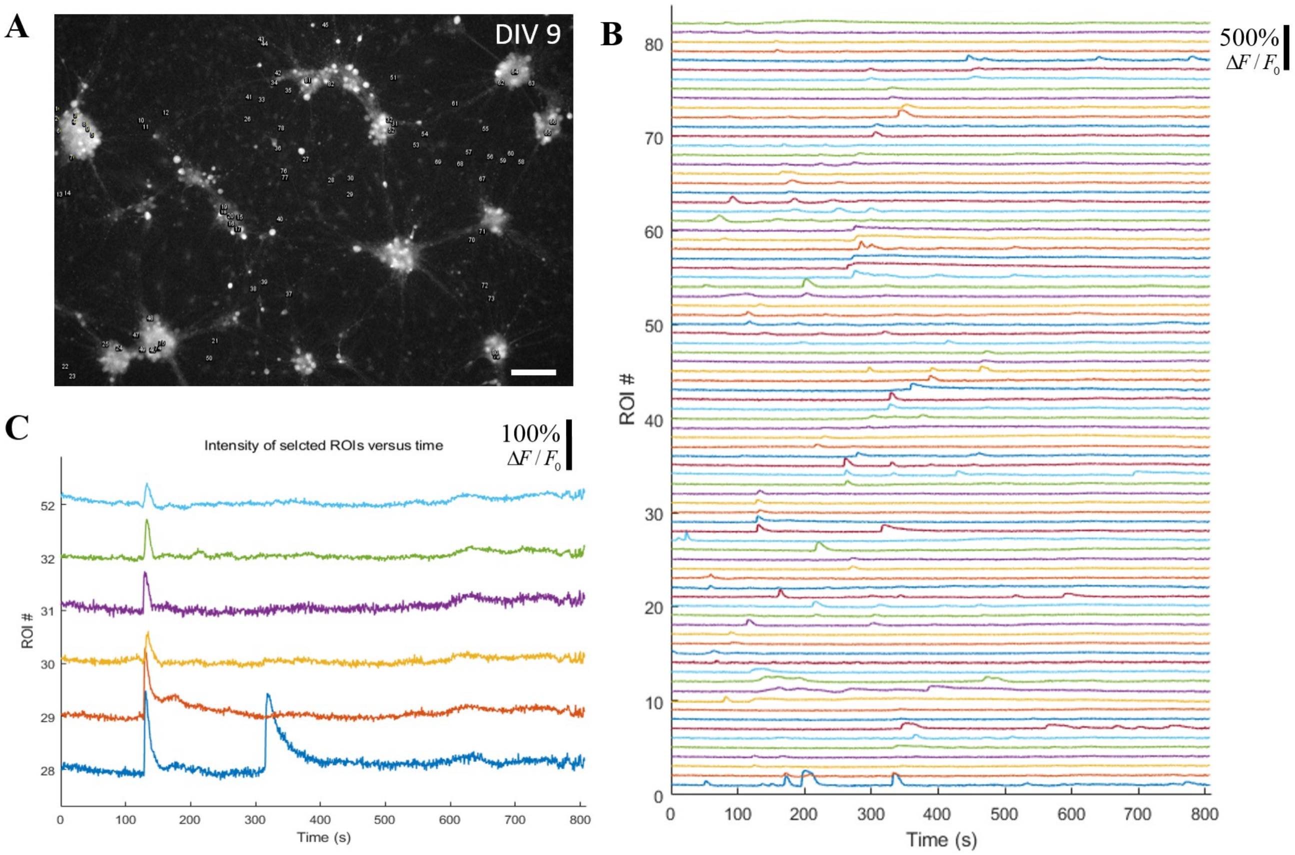


**Fig. S3. Only subnetwork-wide burst activities were observed in a small-cluster large-scale network on day 9.**

(**A**) ROI location in a small-cluster large-scale network on day 9. The grey image is the abstracted green channel of a peak-intensity frame of a calcium recording video. A total of 82 ROIs were selected manually. (**B**) Corresponding traces of relative fluorescence intensity of distinct ROI showed no network bursts, but only sporadic subnetwork bursts. (**C**) A six-neuron subnetwork synchronized calcium elevation on t = 140 s. Scale bar: 100 μm.

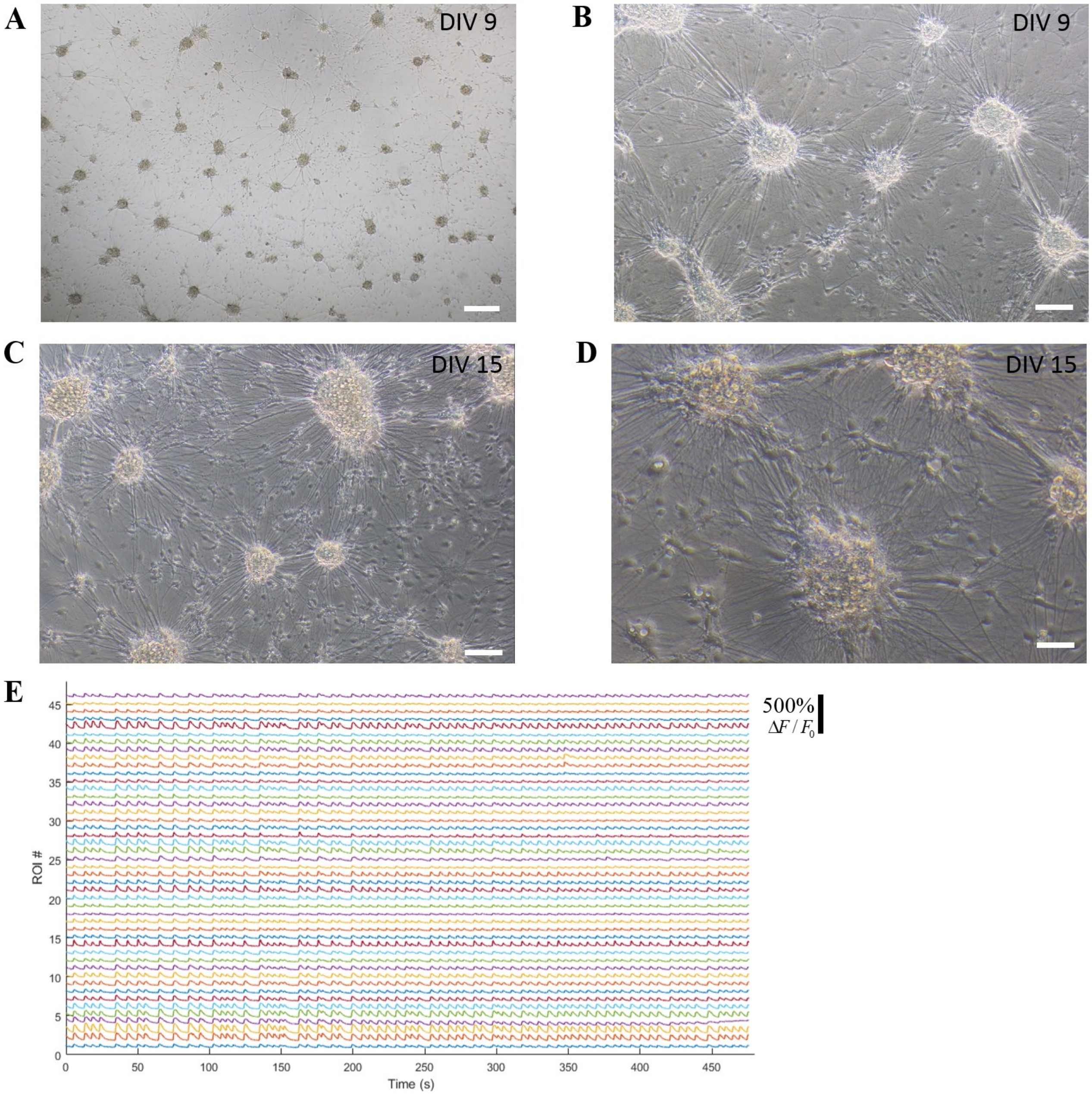


**Fig. S4. Network-wide burst activities and dense neurites were observed in a small-cluster large-scale network on day 15.**

(**A**) and (**B**) Phase-contrast micrographs of the small-cluster large-scale network on DIV 9. (**C**) and (**D**) Phase-contrast micrographs of the small-cluster network on DIV 15 revealed the formation of dense neurites. (E) Relative fluorescence of distinct ROIs in a small-cluster network on DIV 15 showed dominant network bursts. Scale bar: 250 μm, 100 μm, 100 μm and 50 μm in (**A**), (**B**), (**C**) and (**D**), respectively.

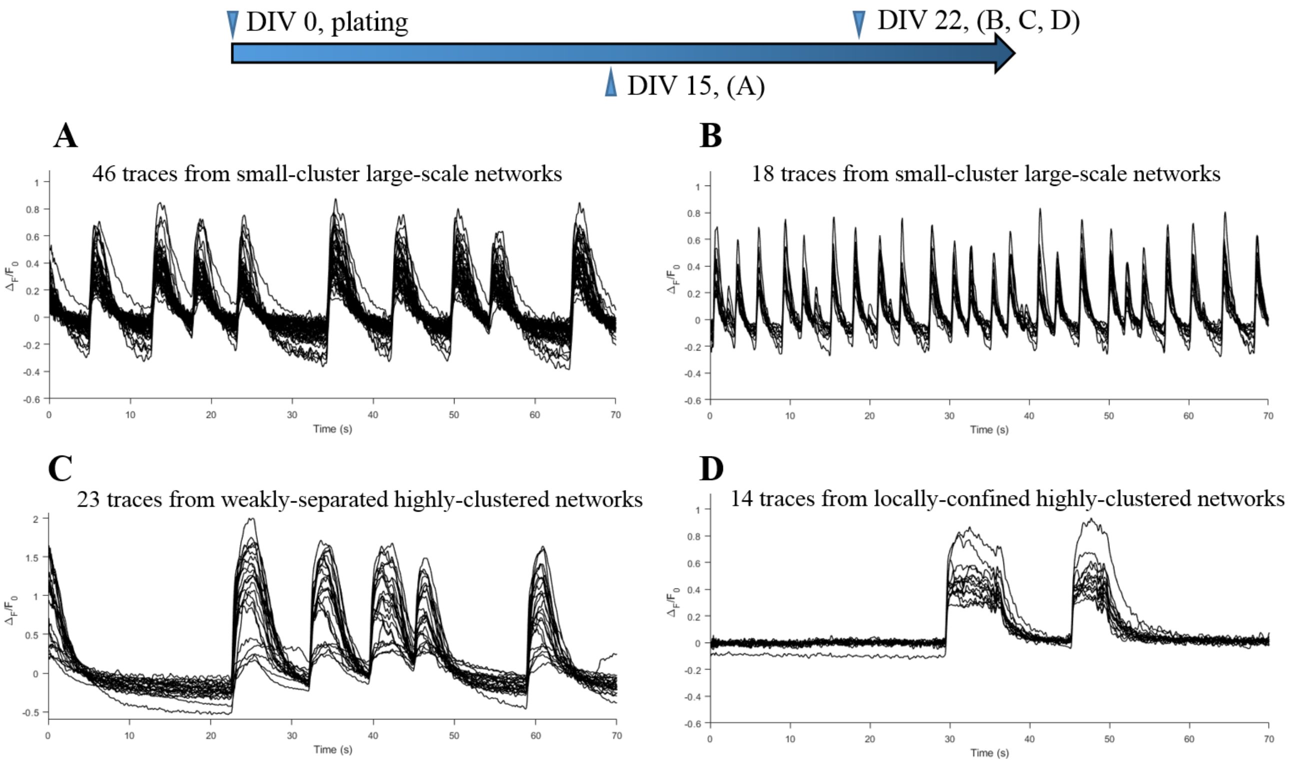


**Fig. S5. Representative overlapped relative fluorescence traces extracted from different cultured cortical networks.**

Traces from small-cluster large-scale networks on day 15 (**A**) and day 22 (**B**) show the high-synchronization of neuronal bursts within a network burst, corresponding to Fig. 2(a) and (b), respectively. Traces from selected synchronized neurons in a weakly-separated medium-scale network (**C**) and from a strongly-separated small-scale network (**D**) also show the synchronization of neuronal bursts within a network burst. (**C**) and (**D**) corresponds to Fig. 4(a) network 1 and Fig. 5(d), respectively.

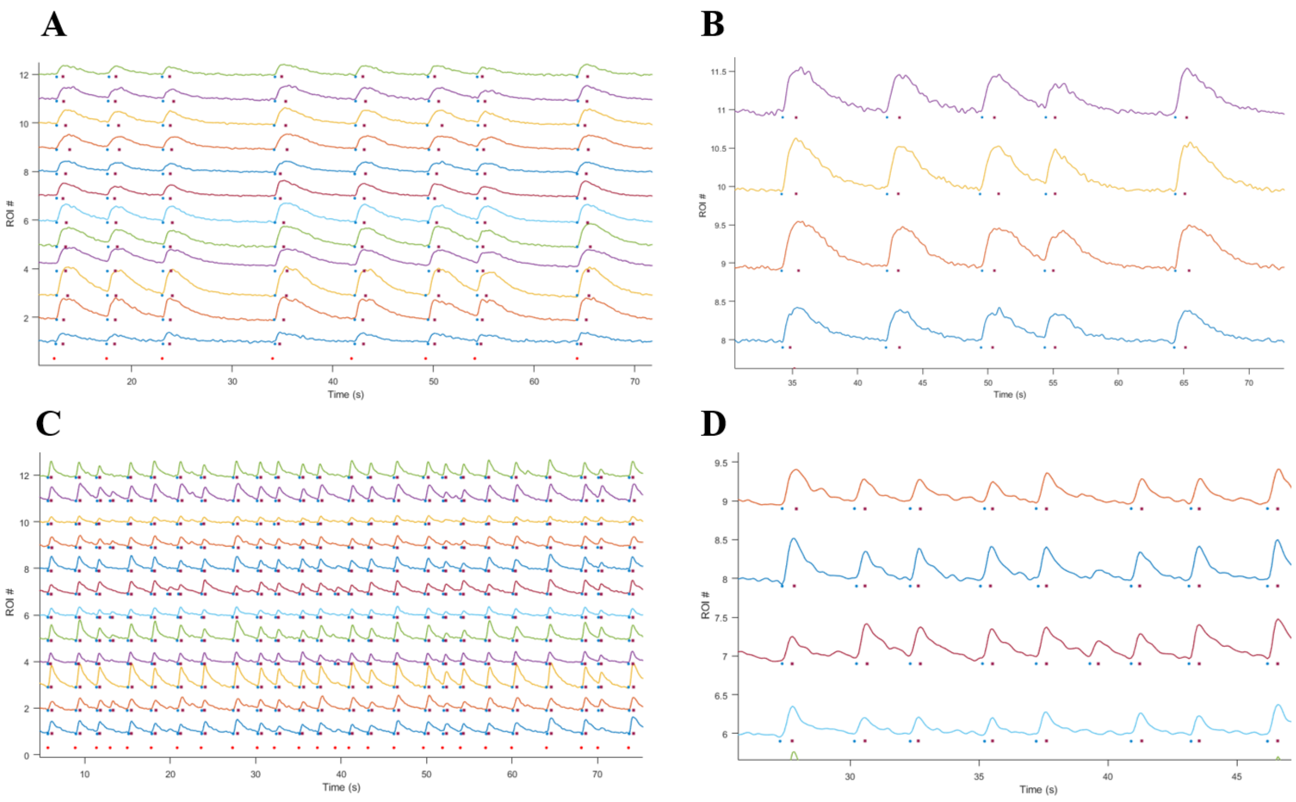

**Fig. S6. Illustration and evaluation of the burst extraction method.**

(**A**) Illustration of one example from a small-cluster large-scale network on day 15. (**B**) Zoom-in view of (**A**). (**C**) Illustration of another example from a small-cluster large-scale network on day 22. (**D**) Zoom-in view of (**C**). Blue round dots, dark-red square dots, and bright-red round dots represent the onset time of a neuronal burst, the end time of a neuronal burst, and the onset time of a subnetwork burst, respectively.

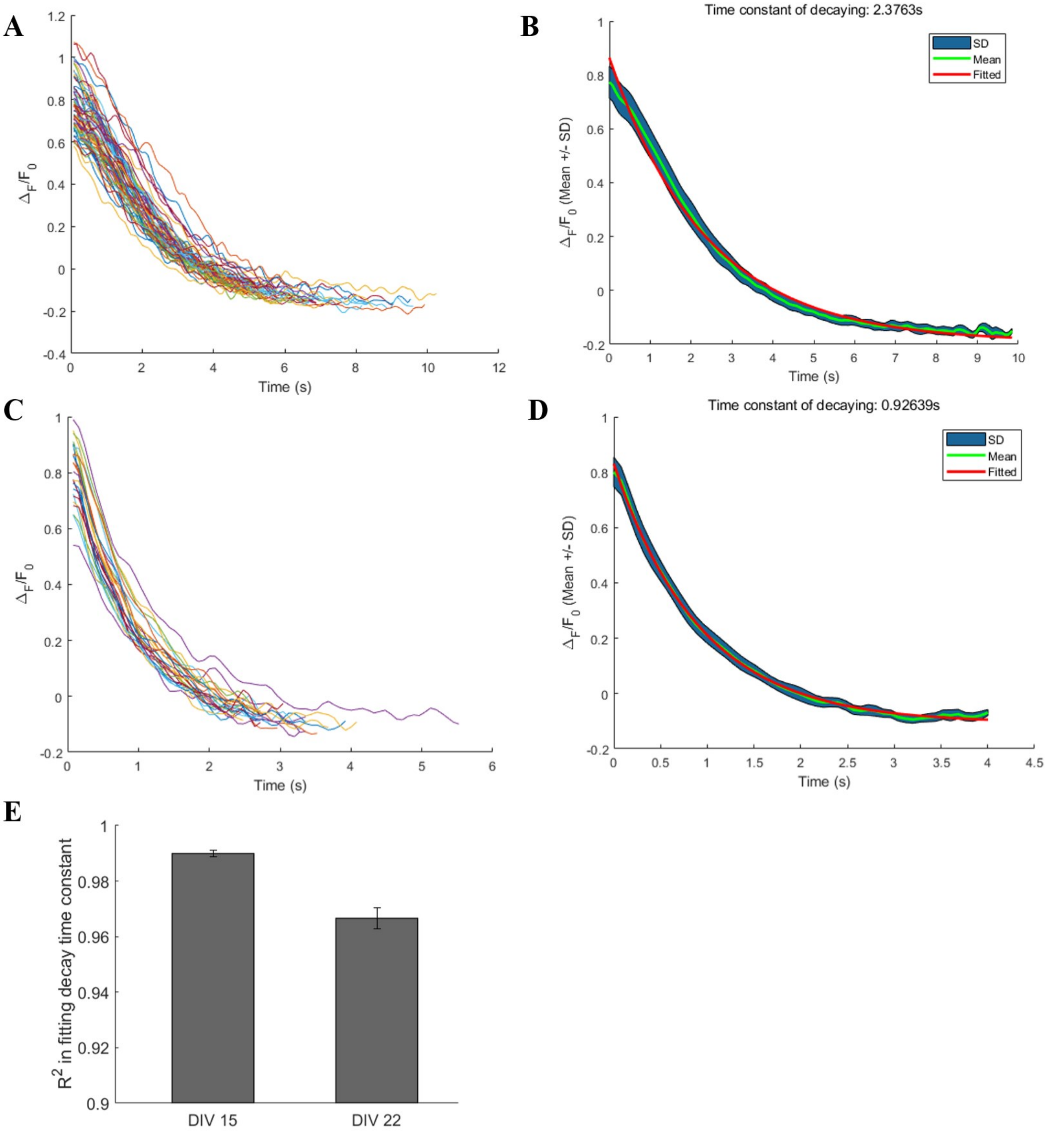


**Fig. S7. Extracting and fitting the calcium decay phase to calculate the decay time constant.** (**A**) and (**C**) Extracted traces of two neurons from small-cluster large-scale networks on day 15 and day 22, respectively. (**B**) and (**D**) show corresponding traces of mean and standard deviation (SD) belt, traces of fitted exponential function, and the decay time constant. The goodness of fit $R^2$ in (**B**) and (**D**) are 0.9909, and 0.9863, respectively. (**E**) $R^2$ in fitting the time constant of decaying phase close to 1 on both day 15 and day 22 demonstrated the feasibility of the choice of exponentially decaying function and the good performance of the proposed method for extracting the decay phase. Data are presented with mean ± SEM.

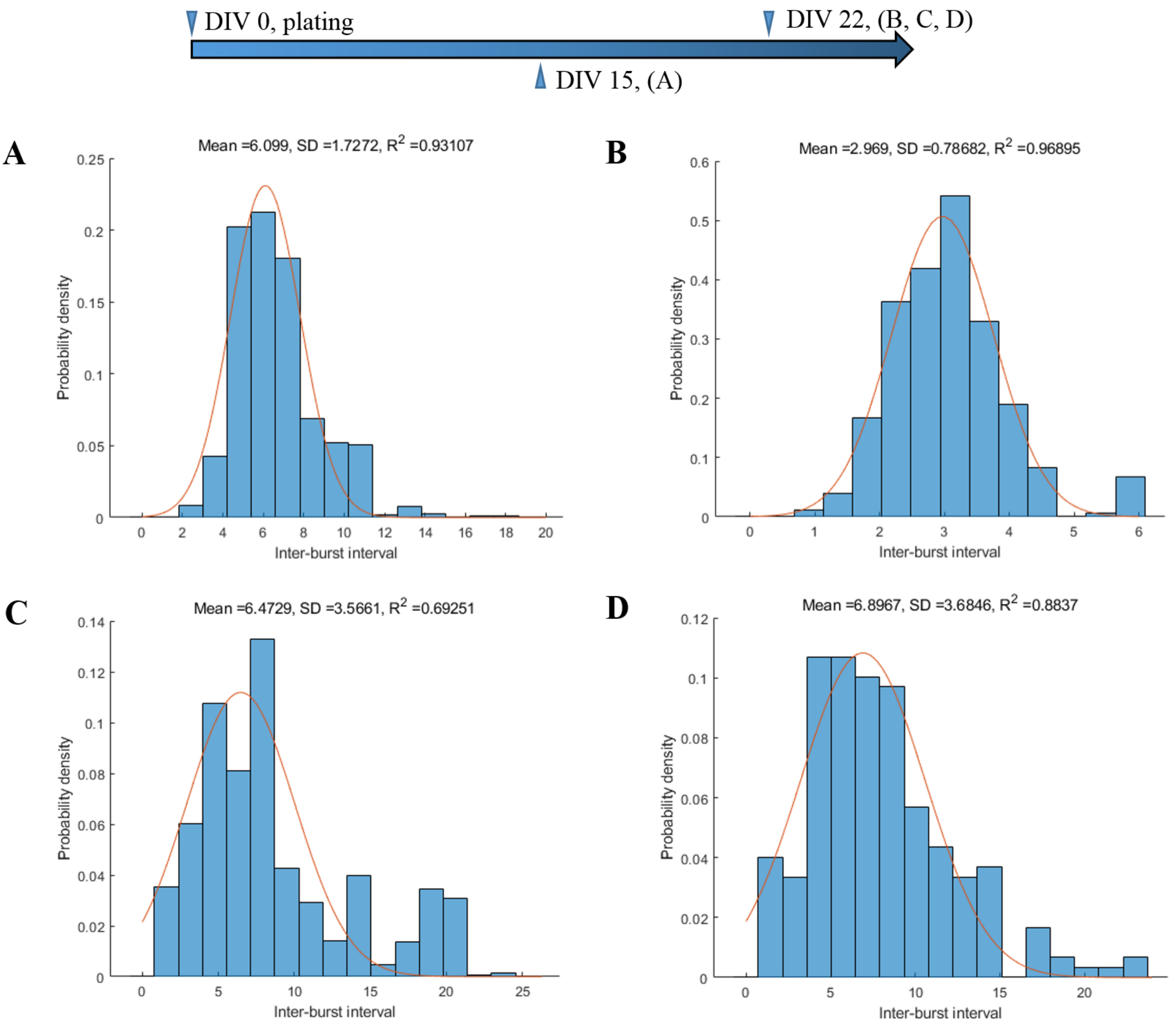


**Fig. S8. The distribution of inter-burst interval (IBI) of different cultured cortical networks.** (**A**) A small-cluster large-scale network on day 15. (**B**) A small-cluster large-scale network on day 22. (**C**) Two weakly-separated highly-clustered networks taken as one merged network on day 22, as in Fig 4(a). (**D**) Another weakly-separated less-clustered network on day 22, as in Fig 4(b). The high values of $R^2$ in (**A**) and (**B**) revealed that the IBI of the small-cluster large-scale network distribute normally. The relatively low value of $R^2$ in (**C**) indicated that the IBI in the merged network distribute non-normally, probably due to the fact that the two subnetworks burst asynchronously. However, in the weakly-separated less-clustered network presenting abundant subnetwork activities, the IBI distribute near normally. The orange curves represent the fitted Gaussian function, with the fitted mean, SD and $R^2$ presented above each subfigure.

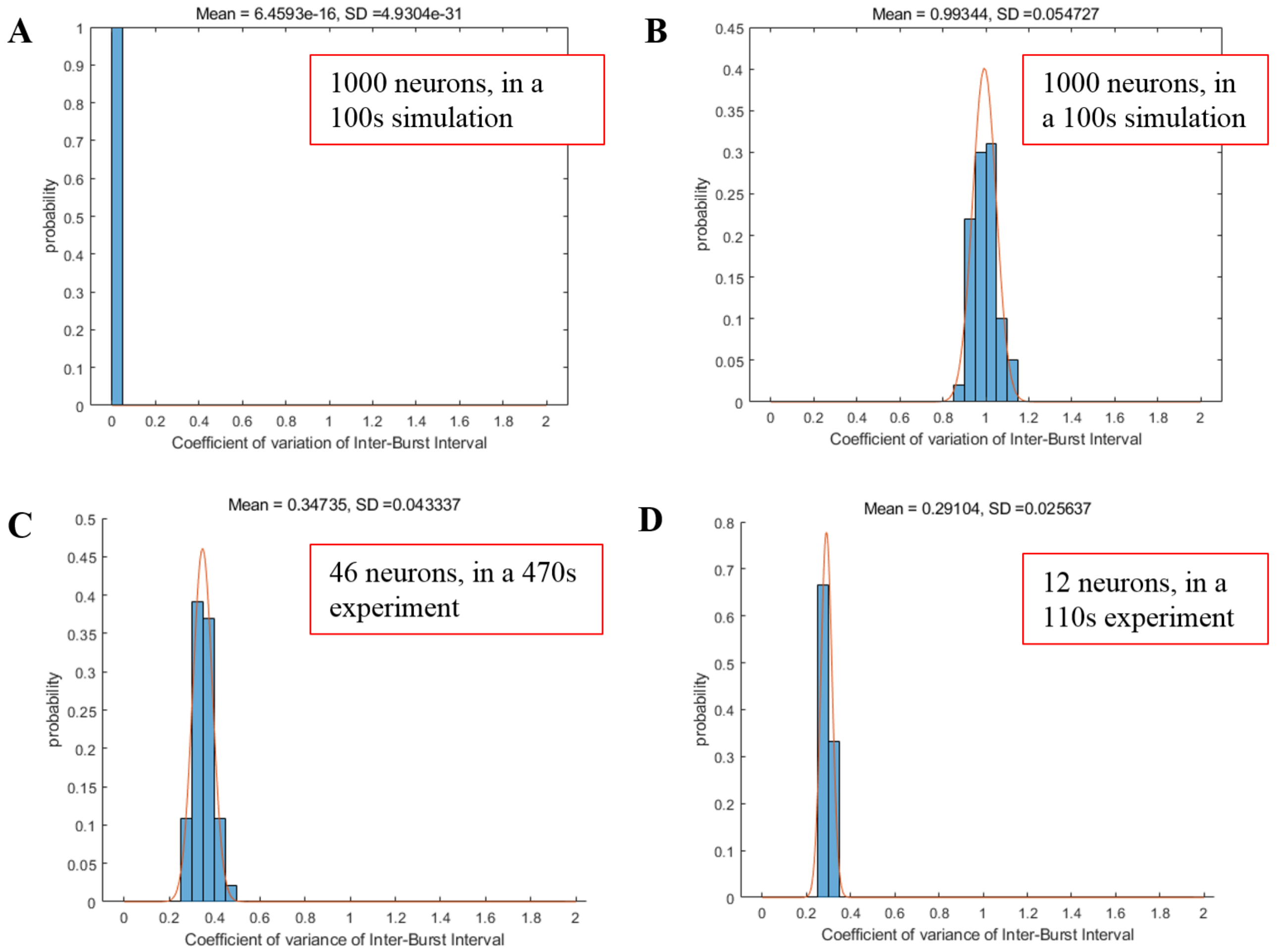


**Fig. S9. The coefficient of variance (CV) of IBI of different cultured cortical networks.**

(**A**) IBICV distribution of a deterministic process in a simulation consists of 1000 neurons with a duration of 100 s. (**B**) IBICV distribution of a Poisson process in a simulation consists of 1000 neurons with a duration of 100 s. (**C**) IBICV distribution of a small-cluster large-scale network on day 15, consisting of 46 recorded neurons with a recording duration of 470 s. (**D**) IBICV distribution of a small-cluster large-scale network on day 22, consisting of 12 recorded neurons with a duration of 110 s. The results revealed that small-cluster networks burst with a combination of a deterministic (rhythmic) process and a Poisson process. The orange curves represent the fitted Gaussian function, with the fitted mean and SD presented above each subfigure.

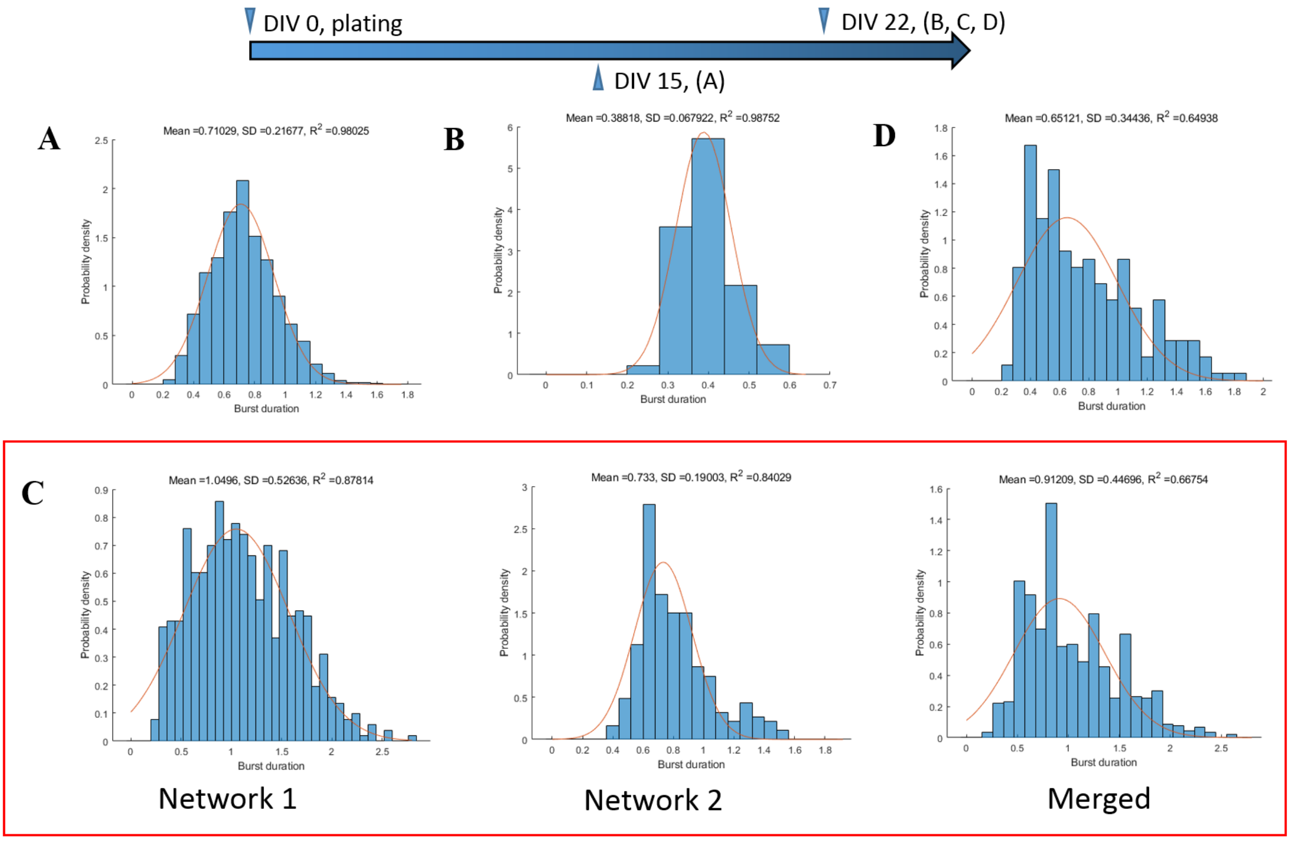


**Fig. S10. The distribution of burst duration of different cultured cortical networks.**
(**A**) A small-cluster large-scale network on day 15. (**B**) A small-cluster network on day 22. (**C**, left) Network 1, (**C**, middle) network 2, and (**C**, right) the merged one from two neighboring weakly separated networks with high clustering degree on day 22, as in Fig 4(a). (**D**) Another weakly separated network with low clustering degree on day 22, as in Fig. 4 (b). The high values of $R^2$ in (**A**) and (**B**) showed that the burst duration in the small-cluster network was distributed normally. The difference of $R^2$ in (**C**) indicated the burst duration of the two networks distributed normally but differently from each other. The low value of $R^2$ in (**D**) suggested that the burst duration of this network was distributed non-normally. The orange curves represent the fitted Gaussian function, with the fitted mean, SD and $R^2$ presented above each subfigure.

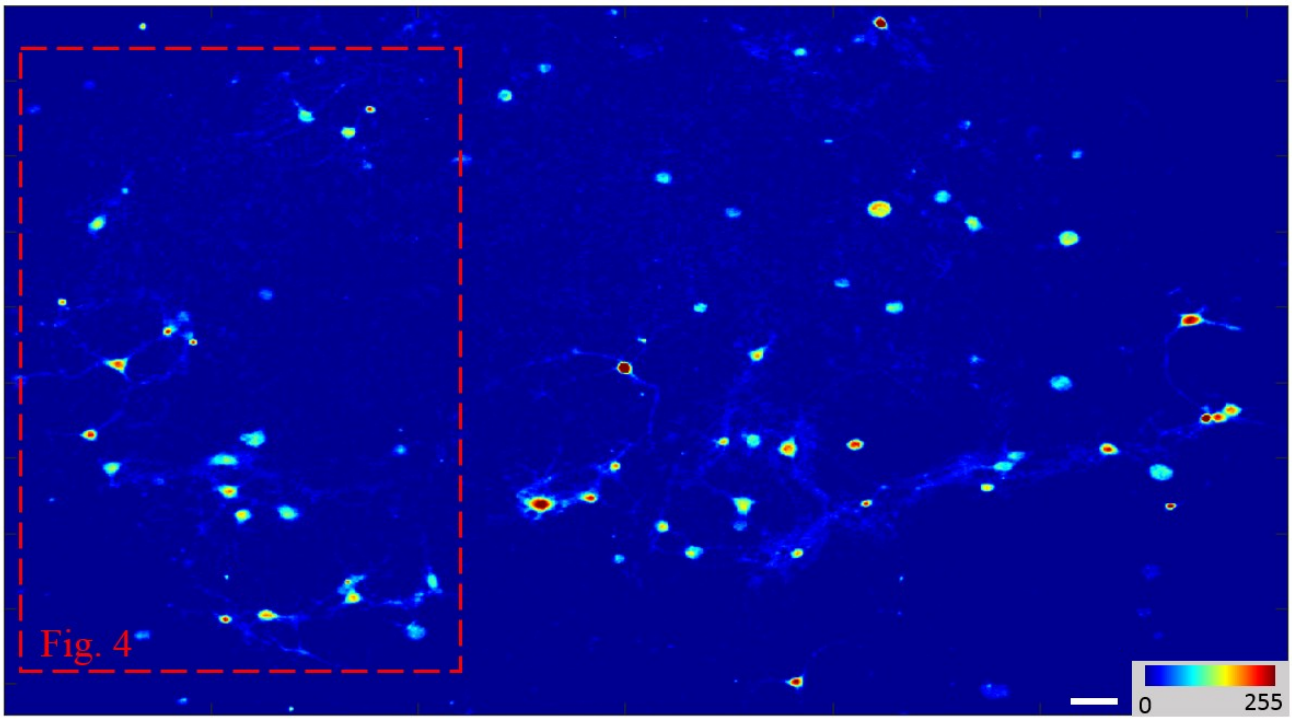


**Fig. S11. Fig. 4 is a subnetwork abstracted from a weakly separated less clustered network.** Scale bar: 40 μm.

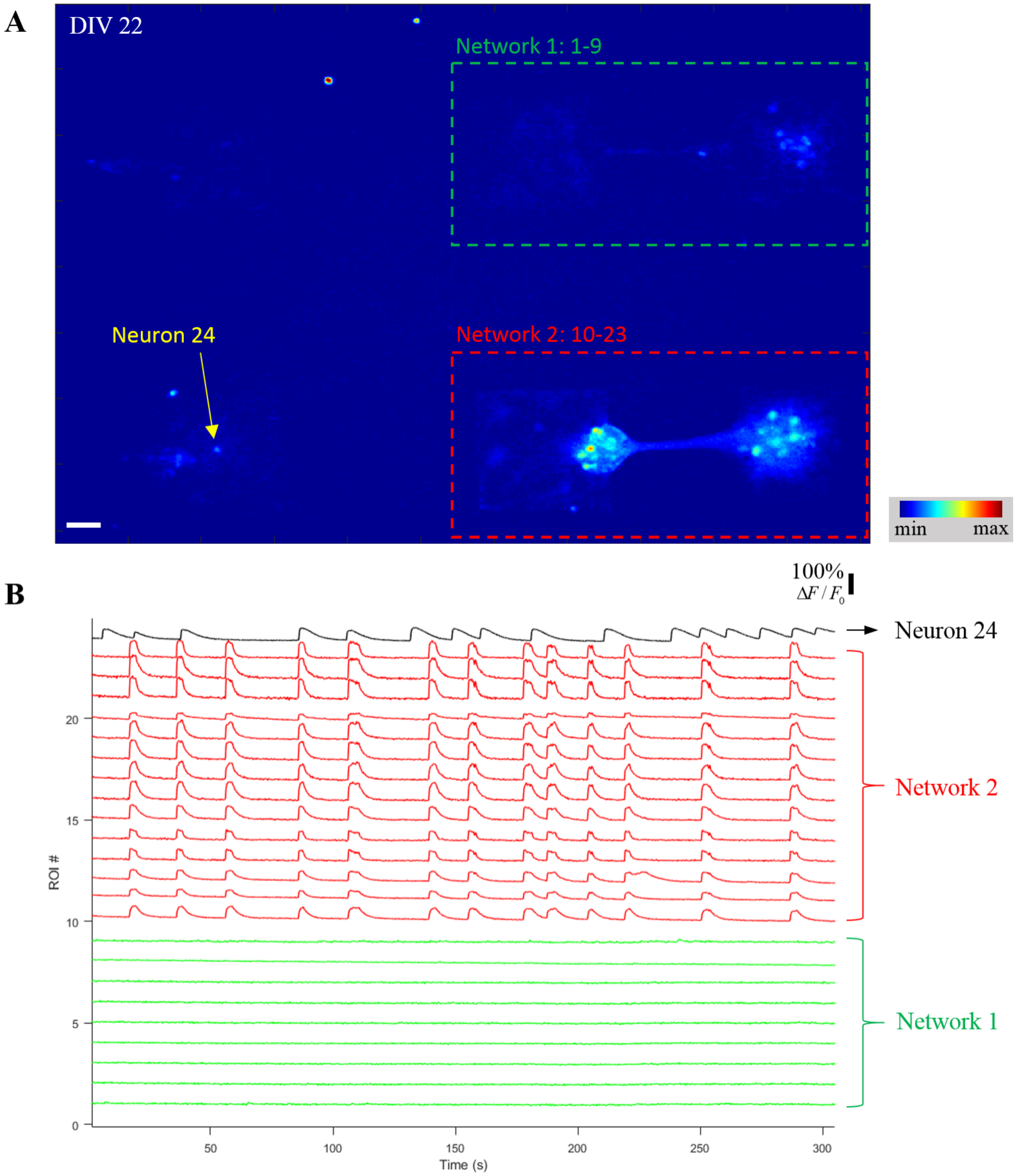


**Fig. S12. An isolated and spontaneously bursting neuron bursts asynchronously with two nearby separated neuronal networks.**

(**A**) False-color fluorescence micrograph of three strongly-separated small-scale networks. (**B**) Corresponding relative-intensity traces on different ROIs from (**A**). The isolated and spontaneously bursting neuron (ROI 24) was labeled with a black trace. Scale bar: 50μm.

### Supplementary Movies

**Movie S1. Synchronized network-wide bursts dominate the calcium dynamics in the small-cluster large-scale network both on DIV 15 and on DIV 22.**

Left, spontaneous calcium activity recording of the Petri dish-wide network on DIV 15, corresponding to Fig. 2(a). Right, spontaneous calcium activity recording of the Petri dish-wide network in another view on DIV 22, corresponding to Fig. 2(b). The small-cluster large-scale network colonizes the entire Petri dish, and the small clusters connect with neighboring ones both structurally and functionally, as evidenced by spontaneous synchronized calcium oscillation. The videos were taken at 12.5 frames/s and replayed at a speed of 10 times compared to real-time. Scale bar: 100μm.

**Movie S2. More complex sub-network calcium activities appear in the less-clustered network induced by sparse seeding.**

Left, spontaneous calcium activity recording of two neighboring highly-clustered networks, corresponding to Fig. 4(a). The two neighboring networks show well-synchronized intra-network bursts and completely out-of-phase inter-network bursts. Right, spontaneous calcium activity recording of another medium-scale network, corresponding to Fig. S11, from which Fig. 4(b) was abstracted. This network is characterized by complex sub-network bursts. The videos were taken at 12.5 frames/s and replayed at a speed of 10 times compared to real-time. Scale bar: 100μm.

**Movie S3. Dominant network bursts reappear in highly-clustered networks induced by local confinement.**

Top left, spontaneous calcium activity recording of an isolated multi-neuron network, where network bursts dominate the calcium dynamics, corresponding to Fig. 5(a). Top right, spontaneous calcium activity recording of an isolated two-neuron network, in which the two neurons show synchronized neuronal bursts, corresponding to Fig. 5(b). Bottom left, spontaneous calcium activity recording of an isolated single-neuron network, in which the isolated neuron present repeated neuronal bursts, corresponding to Fig. 5(c). Bottom right, spontaneous calcium activity recording of a modular 2-by-1 network, which is characterized by a long-lasting post-burst plateau period following each single network burst, corresponding to Fig. 5(d). The videos were taken at 12.5 frames/s and replayed at a speed of 10 times compared to real-time. Scale bar: 50μm.
